# In Silico Identification of Potential Inhibitors of Mycobacterium tuberculosis DNA Gyrase from Phytoconstituents of Indian Medicinal Plants

**DOI:** 10.1101/2024.05.26.596003

**Authors:** Janmejaya Rout, Sayani Das, Sandip Kaledhonkar

**Affiliations:** Department of Biosciences and Bioengineering, Indian Institute of Technology, Bombay, 400076, Maharashtra, India

**Keywords:** DNA gyrase, *Mycobacterium tuberculosis*, natural molecules

## Abstract

DNA gyrase, the sole type-II topoisomerase in *Mycobacterium tuberculosis*, plays a significant role in the bacteria’s survival by catalyzing the DNA topology change and is considered a crucial drug target. The emergence of drug resistance in *Mycobacterium tuberculosis* specifically against the second line of drugs; fluoroquinolones, targeting DNA gyrase, demands new potential compounds that would efficiently interact with it. To find a common suitable inhibitor for the apo and mutated forms of DNA gyrase, in this study, the compounds from an Indian medicinal plant database were screened for selecting potential drug-like molecules and two compounds IM5 and IM7 were selected after successful virtual screening and hit optimization with ADMET predictions, and chosen for MD simulation. The MMPBSA analysis of binding free energy validates the docking results of the two molecules. The principal component analysis also supported the stability of the complexes and thus, these two molecules have turned out to be promising candidates against *Mycobacterium tuberculosis*. Overall, this work sheds light on the potential of DNA gyrase inhibitors as therapeutic agents for *Mycobacterium tuberculosis* treatment, and IM5 and IM7 showed promise as future research compounds.

## 1. Introduction

The emergence of drug-resistant bacterial infections, particularly Mycobacterium Tuberculosis (MTB) causing tuberculosis (TB) poses a significant threat to global public health. The increasing prevalence of multidrug-resistant TB (MDR-TB) and extensively drug-resistant TB (XDR-TB) strains coupled with a lack of novel therapeutic options, [1–3], necessitates the exploration of new treatment strategies. TB has been one of the deadliest bacterial diseases that have remained in society since ancient times [4]. Nowadays due to frequent mutations in bacterial strains, TB patients become resistant to at least one of the first-line drugs such as isoniazid and rifampicin, and are known as Multidrug-resistant TB (MDR-TB) cases. Some TB cases are considered Extensively drug-resistant TB (XDR TB) when the patients show resistance to isoniazid and rifampicin, plus any fluoroquinolone and at least one of the three injectable second-line drugs (*i.e.*, amikacin, kanamycin, capreomycin) [4]. The continuous search for potential antimicrobials to treat these drug-resistant TB cases draws attention to the enzyme DNA gyrase (Gyr), which is a crucial drug target in bacteria. The enzyme DNA Gyr belongs to Type II topoisomerase enzymes, which are widely distributed in the cells and control DNA reorganization. These enzymes act as crucial nucleic acid-dependent nanomachines that regulates DNA topology and govern DNA supercoiling using ATP as a cofactor [5]. Reports have suggested that, the type IIA group of topoisomerase enzymes which are primarily found in bacteria and eukaryotes share sequence homology and conserved evolutionary features in them [6]. Bacteria typically encodes for two types of type IIA topoisomerases which are DNA Gyr and topoisomerase IV. However, in case of *Mycobacterium species,* the enzyme DNA Gyr is the sole type II topoisomerase that takes part in DNA regulation [7]. DNA gyrase serves as the sole target for the clinically relevant class of quinolone derived antibiotics [8] that are used as second line therapeutics in case of mycobacterial infection [9]. During bacterial DNA replication, the enzyme Gyr produces double-stranded DNA breaks to relax the positive super coiling [10]. Thus, the enzyme binds with the DNA and form cleavage complexes. Quinolones bind to these enzyme-DNA complexes and inhibit the replication process, which further leads to chromosome fragmentation followed by cell death [11].

Structurally, the enzyme DNA Gyr is a tetramer composed of two subunits of each GyrA and GyrB. The breakage-reunion domain (N-terminal) and a C-terminal domain form the GyrA subunit while the GyrB subunit consists of the N-terminal ATPase domain and a C-terminal Toprim domain. The DNA segment binds to the breakage-reunion domain of gyrase. When ATP is bound, the N-terminal ATPase domains dimerize and seize the DNA duplex to be transferred. Thus, the subunit A is involved in the DNA binding while the subunit B is associated with the ATPase activity [5,12]. Thus, as a key contributor to bacterial cell survival, the enzyme DNA gyrase could be considered as an ideal target for designing novel therapeutics [6,13].

Several inhibitors of Gyr have been discovered so far, still only the 6-fluoroquinolones class of inhibitors are in clinical use till date [1,14]. This group of inhibitors either act by blocking the ATPase activity or by stabilizing the DNA and Gyr complex preventing successful DNA replication [14]. However, the rise of quinolone resistance due to the mutations in the gene of Gyr in case of multidrug-resistant (MDR) and extensively-drug resistant (XDR) tuberculosis cases is a cause of major concern and needs host-directed therapies for the identification of new treatments [15,16]. Analysis of tuberculosis data from public repositories (NIH) suggests a high prevalence of mutations in the DNA gyrase (Gyr) gene among Mycobacterium tuberculosis strains isolated from patients with MDR-TB and XDR-TB (https://tbportals.niaid.nih.gov/). Primarily, the quinolone resistance-determining regions (QRDR) found in the breakage-reunion domain of GyrA subunit are the sites of mutation that provide bacteria resistance against quinolones [17]. The most common mutations observed in the QRDR region of GyrA are D94G and A90V [17] (https://tbportals.niaid.nih.gov/). Though there are active inhibitors, they have solubility and toxicity issues [18].

In light of the alarming rise of MDR and XDR strains with limited available therapeutic options, there is a critical need for streamlined drug evaluation process and the investigation of abbreviated treatment durations [16]. In that regard, the phytochemicals from medicinal plants have been used for a long to treat human illness [19] could be tested for their potential use. A lot of effort has been made to identify potential phytochemicals to find new physiologically meaningful compounds as therapeutics [20,21]. Moreover, some of the phytochemicals and natural products having specific chemical constituents that show certain biological relevance on various species to achieve high fitness [20,22]. The phytochemicals and natural products have a significant role in discovering and creating therapeutics to treat multiple illness. A recent review article indicates that 34 % of approved small molecule inhibitors over the past 39 years are either natural products or their derivatives [23]. Thus, finding a drug with a pre-validated target like DNA Gyr might be appealing to deal with bacterial resistance as it is absent in humans and functions only in prokaryotic cells like bacteria. The experimental method of finding therapeutics against bacterial infections can be expensive and time-consuming, while computer-aided drug design is a convenient and cost-effective approach widely used for novel drug discovery. One of the popular computational methods to identify novel drugs from large databases is virtual screening. Here, we have implemented well-established computational techniques to identify novel inhibitors of Gyr. First, the Indian Medicinal Plants, Phytochemistry and Therapeutics (IMPPAT 2.0) database [24] containing 17967 phytochemicals was scanned with a drug property filter followed by molecular docking and pharmacokinetic property analysis. Then, the selected compounds were subjected to molecular dynamics simulations to evaluate their stability and conformational dynamics in the binding pocket.

## 2. Materials and Methods

The screening of small molecules, given their drug-like properties using virtual screening, has evolved as one of the most promising techniques in structure-based drug discovery. Several studies report successful drug design applications using docking and MD simulation. This study adopted different methods to screen a small molecule library to identify potent hit compounds based on drug-like properties, pharmacokinetic properties, molecular docking, and MD simulation. The workflow of this study is presented in the **Figure S1**.

### 2.1. Structure preparation of receptor and ligands

The crystal structure of Mycobacterium tuberculosis in complex with MES (PDB ID: 6GAV) for the quinolone resistance determining region (QRDR) was obtained from the RCSB protein data bank [25]. The same structure in complex with AMPPNP (ANP) with PDB ID: 6GAU for the ATP binding site was also obtained from the protein data bank [25]. Both structures are in dimeric form, having full-length sequence similarity. The structures contain GyrA, GyrB with the interdomain connecting region. The only difference between the two structures is the co-crystallized ligand in the two binding sites. We selected chain A from the complex dimers (which contains GyrA, GyrB with the interdomain connecting regions and their co-crystallized ligands) and processed the structures for docking study. The receptor structures were prepared by removing the crystallized ligands and water molecules. Then, the structures were saved in pdbqt format after adding hydrogen atoms and Gasteiger charges in AutoDock tools [26]. The sequence comparison of the PDB 6GAV with the sequences of GyrA and GyrB of *M. tuberculosis* and Escherichia coli (E. coli) obtained from the UniProt was made (**Figure S2**). The mutated structure of the DNA gyrase was made by mutating the residues D747G (D94G) and A743V (A90V) in the QRDR region using Chimera [27]. The ligand structures were obtained from the IMPPAT 2.0 (Indian Medicinal Plants, Phytochemistry and Therapeutics) database [24]. The database currently contains 17967 phytochemicals, including cheminformatics tools to calculate their physicochemical and drug-like properties. The 3D pdb files of these molecules were obtained from the database. The structures were processed by adding hydrogen atoms and Gasteiger charges and then saved to pdbqt format using Open Babel [28].

### 2.2. Binding Pocket Definition and Molecular Docking

Molecular docking was performed on the processed receptor and ligand structures using AutoDock Vina [29]. It uses a hybrid scoring function to estimate the highest affinity conformer of a compound. The crystallized ligand binding sites were considered as the binding pocket for the docking study. A grid box of 30lJÅlJ×lJ30lJÅlJ×lJ30lJÅ size was prepared to cover the binding pockets for all three structures with the centers of the grid boxes set at (70.138, 4.471, 253.971), (79.092, −20.679, 304.253), and (79.092, −20.679, 304.253) for ATP, QRDR, and mutated QRDR site, respectively. The number of binding modes was set to 10, and the exhaustiveness of the search was set to 16 for all the structures. Once the docking run was complete, the docked poses were graded according to their binding affinities. A cutoff value of −7.2 was set to choose the lowest energy docked pose of only those complexes with binding affinity greater than the cutoff. From the docking study, 37 molecules were found above the cutoff limit.

### 2.3. Hit optimization

Some molecules might interact non-specifically with numerous proteins, producing false positive responses during high-throughput screening. These molecules are called pan assay interface (PAINS) compounds. The screened molecules from the docking study were fed to the PAINS remover server (https://www.cbligand.org/PAINS/search_struct.php) to remove the PAINS compounds [30]. The compounds that qualified for the PAINS test were subjected to pharmacokinetic property calculation. The evaluation of physicochemical and pharmacokinetic features of molecules is crucial not only for the initial identification of drug molecules but also for further approval processes during clinical trials. A molecule’s pharmacokinetic property is closely related to its chemical structure. If a molecule in its bioactive form reaches the target site in sufficient concentration to perform the biological event, it is called a potent drug. The assessment of pharmacokinetic properties such as absorption, distribution, metabolism, excretion, and toxicity (ADMET) properties are a crucial factor in the drug development process. The pharmacokinetics and medicinal chemistry properties were explored using the swissADME server (www.swissadme.ch/index.php) [31]. The toxicity parameters of the compounds, such as mutagenic and tumorigenic properties, were also assessed using OSIRIS property explorer [32].

### 2.4. Molecular dynamics

The interaction dynamics of Gyr for the apo and mutated forms in the QRDR site and ATP binding site were monitored using GROMACS-2019 [33]. All the systems were prepared using the CHARMM36 [34] forcefield for the simulations. The small molecule topology parameters were generated using CGenFF [35]. Then, the structures were placed in a dodecahedron simulation box with a spacing of 1 nm from the solute and filled with solvent using the TIP3P water model [36]. The electrical neutrality of the systems was maintained by adding counter ions. The salt concentration of the systems was kept at 0.15 M. The prepared systems were subjected to 50,000 energy minimization steps using the steepest-descent algorithm. After the energy minimization, the systems were set to equilibrium for 100 ps and 1 ns under NVT and NPT ensemble conditions, respectively, at 300 K temperature and 1 atm pressure. A final production run of 100 ns was performed for all the systems with the NPT ensemble. The particle mesh Ewald [37] and force-switching methods were implemented for electrostatic and van der Waals interactions. The velocity-rescaling scheme from the Berendsen thermostat [38] and the isotropic-rescaling scheme from the Parrinello-Rahman barostat [39] maintained the temperature and pressure during the simulation. The time step of the simulation was set to 2 fs, and the frames from the trajectory were extracted at 10 ps intervals. Analysis of the simulation trajectories was made using the Gromacs analysis tools and VMD [40]. The data extracted from the Gromacs analysis tool was plotted using origin 9.0 for further analysis. One of the most popularly adopted techniques, such as molecular mechanics Poisson-Boltzmann surface area (MMPBSA) method, was used from the gmx_MMPBSA tool to find the binding free energy of the systems [41]. The binding free energy was calculated from the MD trajectories, taking 1000 snapshots from 90-100 ns regions. The results were extracted to DAT and CSV files and analyzed with the gmx_MMPBSA_ana module.

### 2.5. Principal component analysis

Principal component analysis (PCA) is a popular multivariate statistical method that can assign a new set of variables to the structure in each frame of a MD trajectory with little information loss, known as principal components (PCs) [42]. Here we employed the g_covar module from Gromacs to calculate and diagonalize the covariance matrix giving the eigen values that are the PCs corresponding protein conformations from the MD trajectory [43]. The structural data was represented in a lower dimension by projecting the distribution to subspace defining largest PCs with g_anaeig tool. Their corresponding eigenvector values describe the fraction of the overall variance of atomic coordinate changes registered in each dimension. Typically, in PCA, only a small number of dimensions are required to roughly capture 70% of the variance in the structures under study [44]. As a result, while preserving most of the variance in the original distribution, the first few eigenvectors are adequate to offer a meaningful description [45]. The g_sham module was utilized to get the free energy landscape plot from the top two eigenvectors obtained from the PCA.

## 3. Results and Discussion

### 3.1. Identification of potential drug-like molecules

Estimating drug-like properties is one of the most critical parameters, as it saves time and effort at later drug testing stages. The drug-like properties of a chemical compound would help determine the ability of a compound to be used as an orally administrable drug for humans. Here, the library of compounds was screened following different drug-like filters such as Lipinski [46], Ghose [47], GSK 4/400 [48], Pfizer 3/75 [49], Veber [50], and Egan rule [51]. The molecules with zero violations for the above rules and a quantitative estimate of drug-likeness (QEDw) score greater than 0.8 were selected for further studies. From 17967 molecules of IMPPAT 2.0, 104 molecules qualified for drug-like properties.

### 3.2. Binding site information and Virtual screening

Active form of DNA gyrase is a heterotetrameric enzyme having GyrA and GyrB and both units play a crucial role in the DNA cleavage and relegation [5]. Here, we have taken a full-length structure of gyrase that consists of GyrA, GyrB, and interdomain connections. In this full-length structure, the GyrB ligand binding site is called the ATP binding site, and the ligand binding in the GyrA QRDR region is referred to as the QRDR site in this study. The molecular electrostatic potential (MEP) energy surface plots (**Figure S3**) for the respective reference compounds, ANP and MES in the ATP and QRDR sites were obtained using PyMol [52] for the binding pocket information. Plots of MEP for IM5 and IM7 (details are provided in ***Section 3.4***) in the ATP, QRDR, and mutated QRDR sites are also provided in **Figure S3**. The MEP surface analysis can provide information on the macromolecule’s active site, indicating the relative ligand orientation and active site nature where an approaching electrophile could interact [42]. From the plot, it is observed that the co-crystalized ligand ANP binds in a highly electronegative region of the binding pocket in the ATP binding site (**Figure S3(a)**) while MES in the QRDR site accommodates itself in a less electropositive region (**Figure S3(b)**). This implies that both the co-crystallized ligands were stabilized by polar interactions in the active site.

Structure-based virtual screening was performed in the ATP binding site, QRDR, and mutated QRDR site on the drug-like filtered compounds to find suitable inhibitors. A library of natural molecules (IMPPAT 2.0) was selected for the structure-based screening to find out potential inhibitors of DNA gyrase [24]. The virtual screening aims to find a common candidate which could effectively act on both domains of Gyr, i.e., ATP binding site and QRDR site. The search volume around the binding pocket was defined considering the binding site of the co-crystallized ligands in the obtained PDB files so that the docking algorithm acts effectively within the search volume. A threshold value of −7.2 kcal/mol was set to filter out a common potential inhibitor for both drug target sites. With the specified cutoff, 37 compounds were selected as hit compounds with their lowest energy docked pose as the best conformation.

### 3.3. Optimization of the hit compounds

Selected compounds from the virtual screening were subjected to a false positive remover program (https://www.cbligand.org/PAINS/login.php) to filter out the PAINS (pan assay interference) molecules. Among all the molecules, 36 molecules qualified PAINS test. Then, these molecules were analyzed with the OSIRIS property explorer to evaluate their drug-toxicity parameters (**Table S1**). This analysis excluded three compounds, two of which were tumorigenic and one mutagenic. The remaining 33 compounds were subjected to swissADME (http://www.swissadme.ch/) for the pharmacokinetic property analysis (**Table S2**). Fourteen compounds were found to be positive substrates for P-glycoprotein, which led to their exclusion from the lead library. Then, from the remaining 19 molecules, only two were observed to be the non-inhibitors of any of the cytochrome P450 isoforms, while the remaining 17 were the inhibitors of at least one of the CYP isoforms (**Table S2**). So, only these two compounds were selected as lead compounds and considered for stability analysis of their complexes using MD simulation.

### 3.4. Docking analysis of the selected lead compounds

The co-crystallized ligands and the hit compounds were docked in the receptor’s binding sites. The complexes were analyzed with respect to binding affinity, number of hydrogen bonds, and nature of interacting residues. The reference compound ANP showed a binding affinity of −7.2 kcal/mol for the ATP binding site, while the co-crystallized ligand (MES) in the QRDR and mutated QRDR site showed an affinity of −5.0 kcal/mol and −5.1 kcal/mol, respectively. Considering the cutoff value of −7.2 kcal/mol, 37 molecules (**Table S1**) were selected, among which IMPHY006072 showed the highest affinity in the ATP binding site with a binding affinity of −8.6 kcal/mol. In contrast, the highest binding affinity of −9.8 kcal/mol was observed for IMPHY003954 in the QRDR and −9.6 kcal/mol for IMPHY003529 in the mutated QRDR site. But after the pharmacokinetic property filter (as discussed in ***Section 3.3***), only two molecules were selected: IMPHY005869 (IM5) and IMPHY007403 (IM7). The binding affinity of IM5 was −7.4 kcal/mol for the ATP and QRDR site, while it was −7.3 kcal/mol for the mutated QRDR site. The lowest energy docked score of IM7 was observed to be −7.3 kcal/mol, −7.5 kcal/mol, and −7.6 kcal/mol for ATP, QRDR, and mutated QRDR sites, respectively. The lowest energy docked pose of the respective co-crystallized ligands, IM5, and IM7 in the ATP and QRDR site with their 2D interaction diagram is shown in **Figure 1**. The docked conformations of IM5 and IM7 with the highest binding affinity in the mutated QRDR site is provided in **Figure S4** and **S5**, respectively. The **Figure 1(a)** shows that ANP has hydrogen bonding (H-bond) interaction with H20, R16, and K348 residues in the ATP site. The residues I284, R169, D615, T347, E24, and T349 are associated with hydrophobic interactions with ANP (**Figure 1(a)**). There are four hydrogen bonding residues (H20, R16, K348, and D615) observed in the case of IM5 (**Figure 1(b)**), while IM7 (**Figure 1(c)**) had five hydrogen bonds (H20, R16, K348, R169, and D615) in the ATP binding site of Gyr. Also, other hydrophobic interacting residues were observed in both the ligand complexes (**Figure 1(b, c)**), resulting in the stability of their interactions.

**Figure 1.**
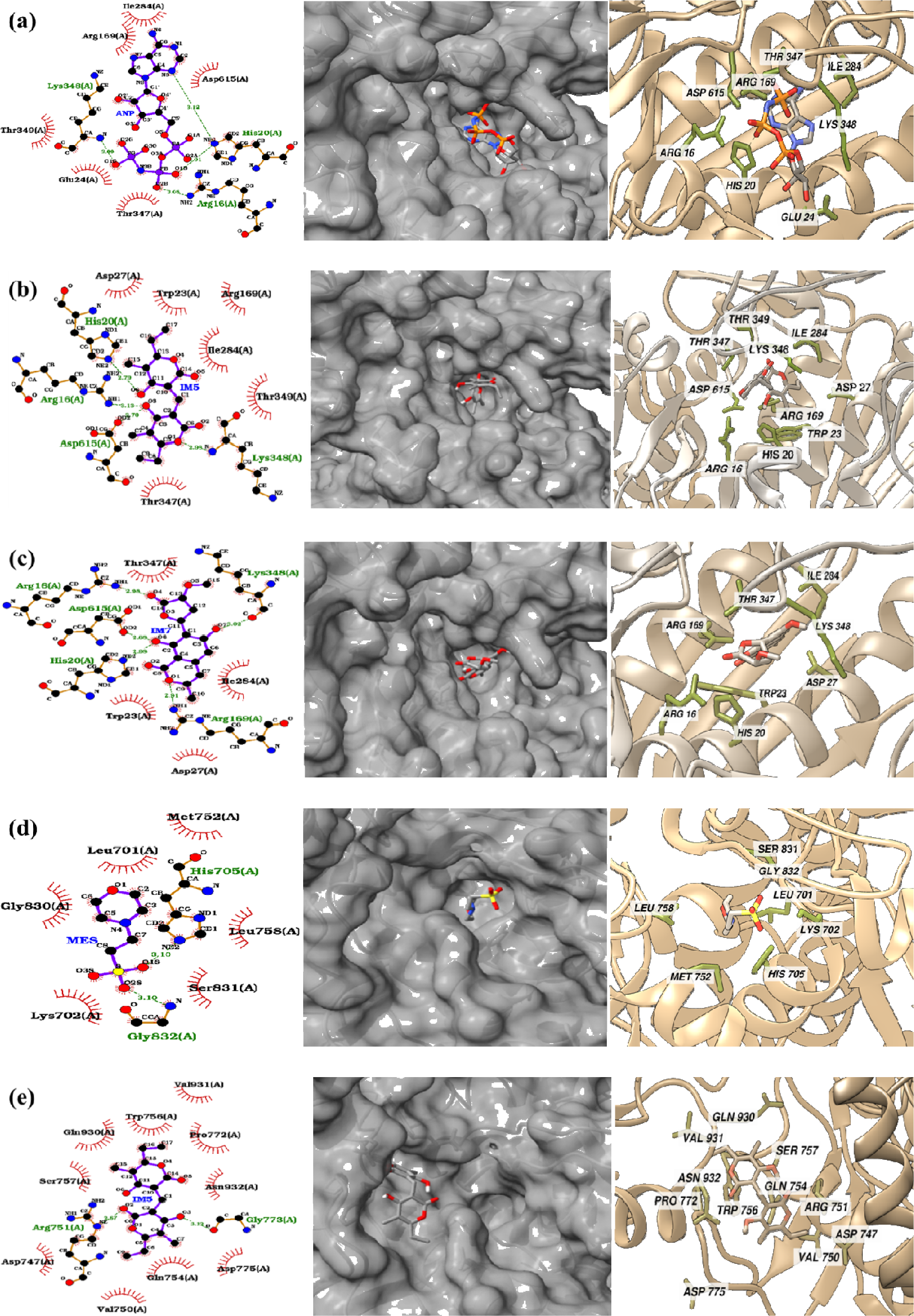
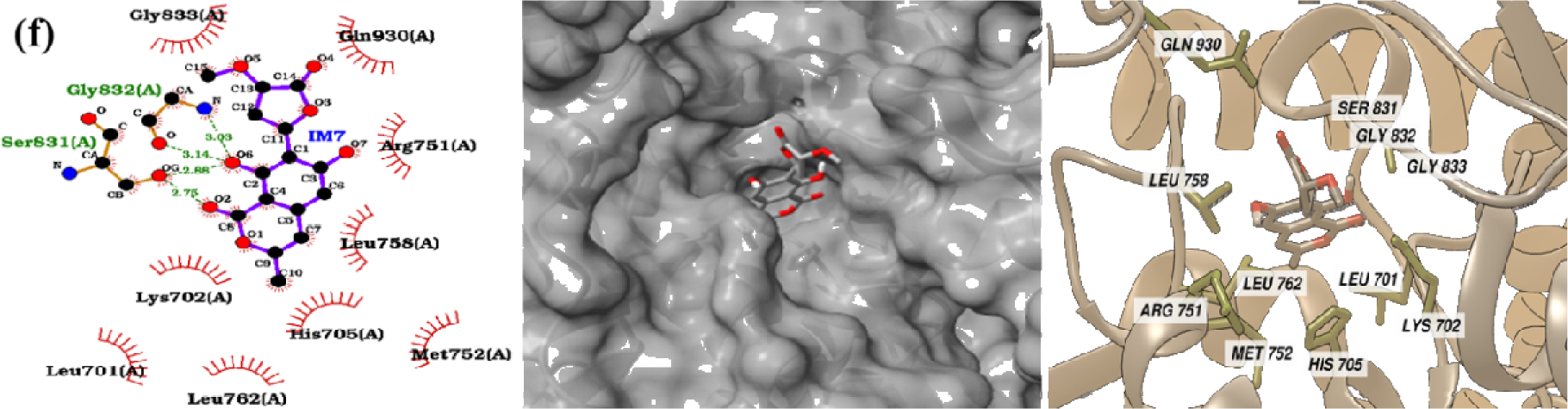
Respective 2D interaction diagram, surface, and 3D binding site view of the lowest energy docked pose of reference (ANP (a)) and lead compounds (IM5 (b), and IM7 (c)), in the ATP binding site. The reference (MES (d)) and lead compounds (IM5 (e), and IM7 (f)), in the QRDR site as obtained from molecular docking and pharmacokinetic filters.

The ligand interactions with Gyr in the QRDR site is provided in **Figure 1(d-f)**. MES (H705 and G832) and IM5 (R751 and G773) had two H-bonds, while IM7 had four H-bonds with two residues (S831 and G832) of Gyr as seen from **Figure 1(d), (e)**, and **(f)**, respectively. Six hydrophobic interactions were observed for MES with Gyr (**Figure 1(d)**), while ten were observed for IM5 (**Figure 1(e)**). Nine hydrophobic interacting residues were found for IM7 in the QRDR site of Gyr (**Figure 1(f)**). We have also performed docking studies for a few possible mutations that are reported [17] for the residues A743 and D747 and are presented in **Table S3**. The lowest energy docked pose with the 2D interaction diagram of IM5 and IM7 in the mutated sites of Gyr are provided in the **Figure S4** and **S5**, respectively. The binding affinity (**Table S3**) of IM5 and IM7 were observed to be almost same for all the mutations performed on A743 and D747. Also, the H-bonding residues are conserved for all the mutations as seen from the docking studies (**Table S3**). In the QRDR site of mutated Gyr, IM5 had two H-bond (R751 and G773) interactions and nine hydrophobic interactions (**Figure S4(j)**) for the most common mutation observed (A743V and D747G). The interaction of IM7 for the same mutation in the QRDR site of Gyr was stabilized by three H-bond interactions with residues S831 and G832 (**Figure S5(j)**). Also, IM7 had nine hydrophobic interactions with the Gyr residues in the same mutated QRDR binding pocket.

The MEP surface analysis of the IM5 and IM7 docked complexes in the ATP, QRDR, and mutated QRDR sites was also performed to monitor the nature of the interaction site (**Figure S3**). In the ATP binding site (**Figure S3(a)**), both the IM5 and IM7 were found in the electronically rich regions of the binding pocket, suggesting their interaction to be in the polar site. In the case of the QRDR (**Figure S3(b)**) and mutated QRDR (**Figure S3(c)**) sites, IM5 interacts with an electron-rich region, while IM7 interacts with an electron-deficient region, suggesting that both interact in potentially active regions, which implies their underlying interaction stability.

### 3.5. Analysis of MD simulation studies

The structural stability and conformational dynamics of the gyrase-ligand systems were monitored using MD simulation. The stability of the docked complexes was analyzed by monitoring the systems’ root-mean-square deviations (RMSD) with respect to their initial conformations from a 100 ns MD simulation. The RMSD plots of Gyr with ligands ANP, IM5, and IM7 in the ATP site are shown in **Figure 2(a)**. The figure shows that all the simulations attained equilibrium within ten ns of the simulation time and remained stable for entire simulation duration of 100 ns. The RMSD value of ANP is 4.05 ± 0.46 Å, while for IM5 and IM7, it was observed to be 3.85 ± 0.52 Å and 5.24 ± 0.83 Å, respectively. The RMSD values imply that IM5 had the least conformational change compared to ANP and IM7 during the simulation. However, the RMSDs show a similar pattern for ANP, IM5, and IM7, which shows their stable nature during the simulation.

**Figure 2.**
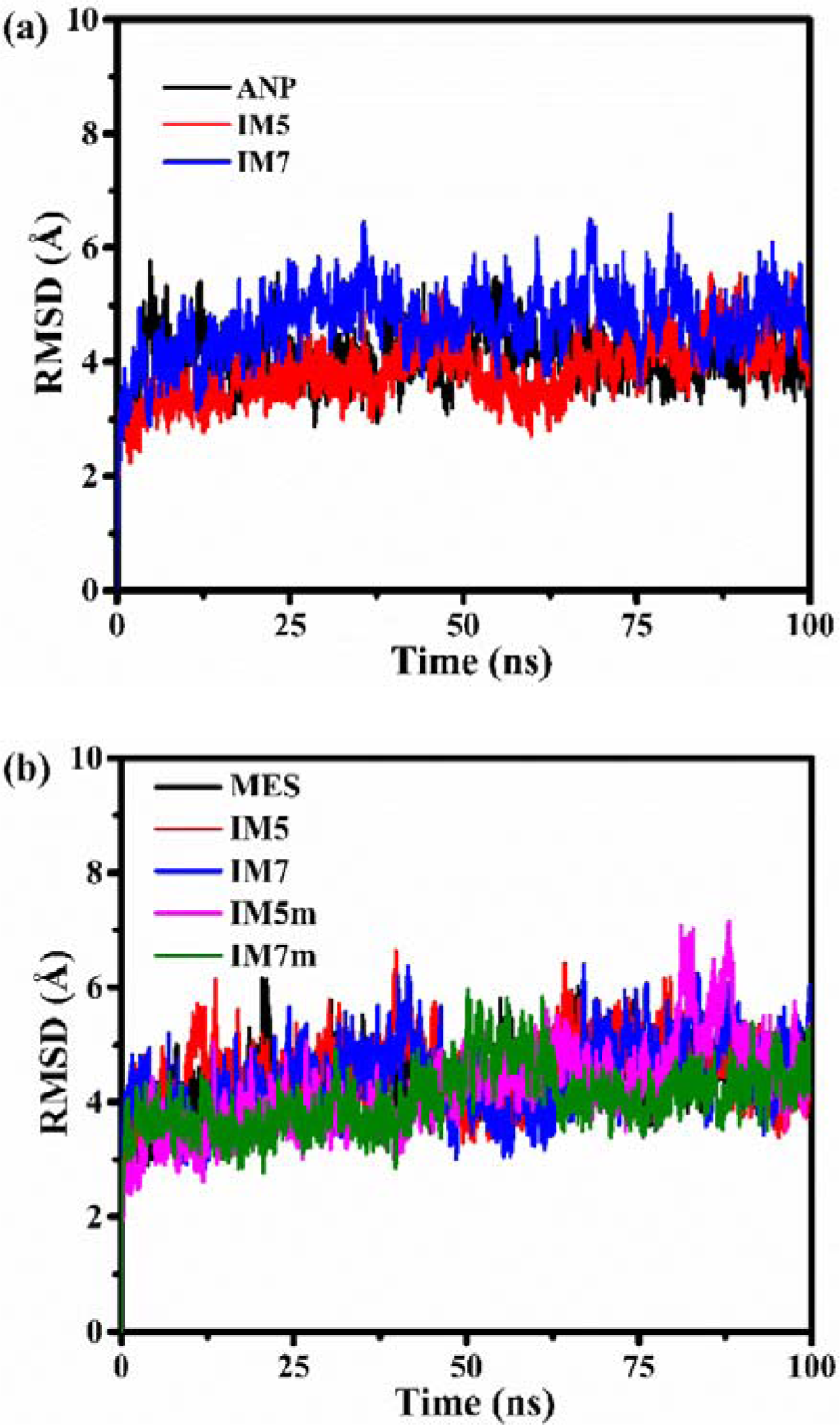
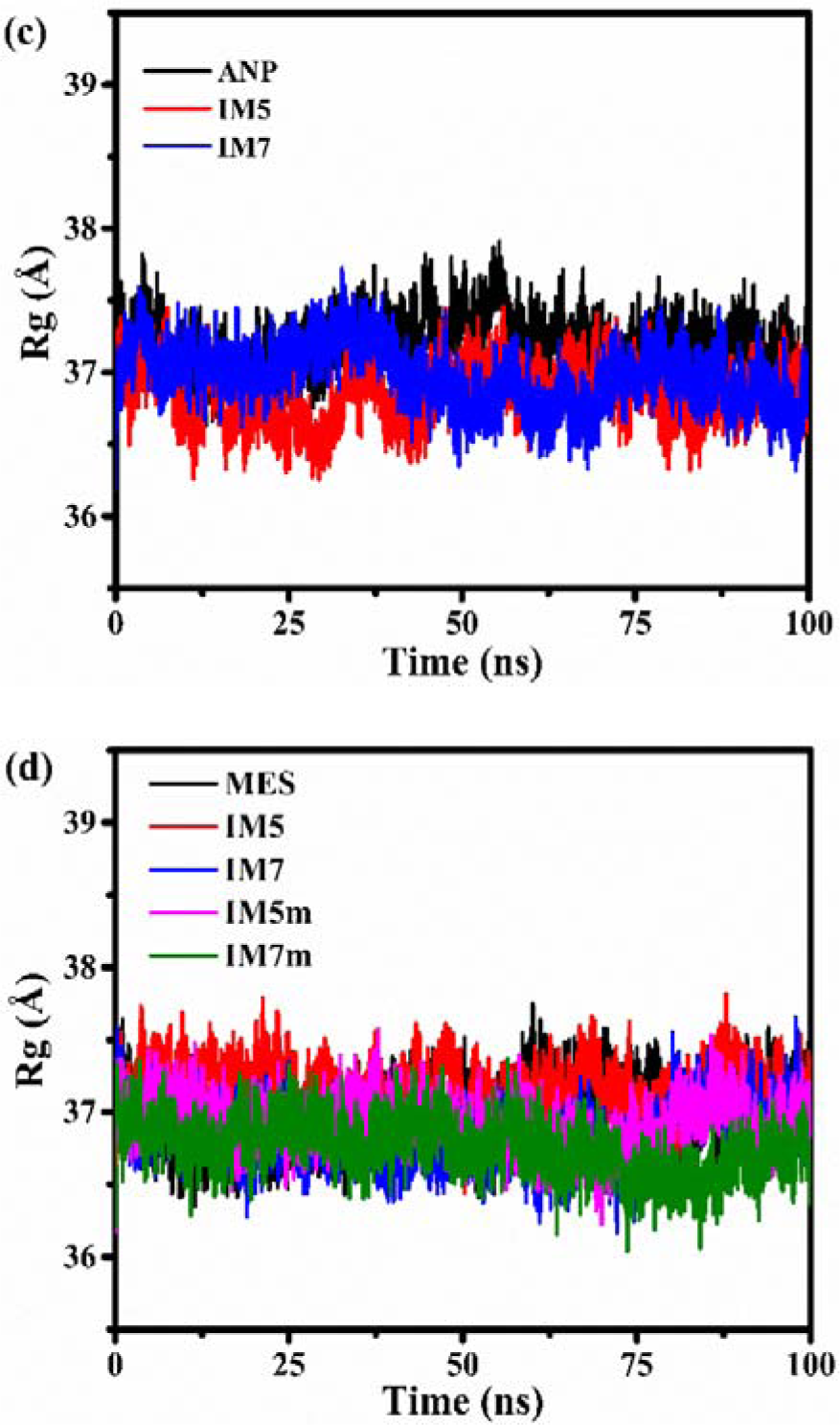
**(a)** Root-mean-square deviation of reference (ANP) and lead (IM5, IM7) compounds in the ATP site; **(b)** Root-mean-square deviation of reference (MES) and lead compounds in the QRDR (IM5, IM7) and mutated QRDR (IM7m, IM7m) site; **(c)** Radius of gyration of reference (ANP) and lead (IM5, IM7) compounds in the ATP site; **(d)** Radius of gyration of reference (MES) and lead compounds in the QRDR (IM5, IM7) and mutated QRDR (IM7m, IM7m) site.

The RMSD plot for MES, IM5, and IM7 in the QRDR site and the mutated QRDR site for IM5 and IM7 is shown in **Figure 2(b)**. All the systems attained equilibrium within 10 ns of the simulation time with similar patterns of fluctuations. The RMSD of MES, IM5, and IM7 was observed to be 4.31 ± 0.54 Å, 4.50 ± 0.58 Å, and 4.42 ± 0.63 Å, respectively, in the QRDR site, while for the mutated QRDR site, the RMSD was 4.21 ± 0.76 Å and 4.04 ± 0.52 Å for IM5 and IM7. The values of RMSD for IM5 and IM7 in the mutated site suggest that these ligands are in a state of more stable conformation that leads to less RMSD value than that of the QRDR site.

The compactness and stability of a protein structure can be gauged through its radius of gyration. **Figure 2** shows the plot of the Rg data of the ATP, QRDR, and mutated QRDR site for the simulation duration. As seen from the figure, IM5 and IM7 showed similar Rg patterns concerning the reference compounds both in the ATP site (**Figure 2(c)**) and QRDR site (**Figure 2(d)**). The Rg plot in the mutated QRDR site (**Figure 2(d)**) is also of a similar nature. The average Rg values calculated for ANP, IM5, and IM7 in the ATP site are 37.19 ± 0.20 Å, 36.84 ± 0.20 Å, and 37.20 ± 0.27 Å, respectively. The values suggest that IM7 bound to Gyr is a bit more compact than the other two, indicating the system’s enhanced stability. In the QRDR site, the Rg values corresponding to MES, IM5, and IM7 are 36.99 ± 0.22 Å, 37.15 ± 0.18 Å, and 36.84 ± 0.19 Å, respectively. Similarly, the Rg values for the mutated (A743V and D747G) QRDR site are 36.93 ± 0.17 Å and 36.75 ± 0.19 Å for IM5 (IM5m) and IM7 (IM7m). In the QRDR site, a higher Rg value of the Gyr-IM5 complex suggests a more compact conformation than the other ligands. Generally, all complexes had a stable structure in the presence of the ligands, as seen from their respective Rg values.

The plot for the root-mean-square fluctuation (RMSF) of Gyr with the lead ligands was provided in **Figure 3**. The RMSF measures the movement in the elemental coordinates of a structure from its reference structure. The plots show that the Gyr structure in the presence of different ligands is stable as it had fewer fluctuations except for the turn and loop regions, which are spiked fluctuations (**Figure 3(a, b)**) in the structure. The residues having highest fluctuations in the turn and loop regions are labelled in **Figure 3(a, b)**. The average RMSF values for ANP, IM5, and IM7 are 1.92 ± 1.01 Å, 1.95 ± 1.28 Å, and 2.73 ± 1.54 Å, respectively, for the ATP site, while it is 2.33 ± 1.23 Å,2.05 ± 1.23 Å, and 1.92 ± 1.10 Å for MES, IM5, and IM7 in QRDR site. The RMSF values suggest that the Gyr-IM7 complex is attaining a flexible structure with respect to the reference structure complex (Gyr-ANP) and Gyr-IM5 complexes. But for the QRDR site, RMSF values are less for Gyr-IM5 and Gyr-IM7 complex than the reference, i.e. Gyr-MES. Also, fewer RMSF values were observed for IM5m (1.96 ± 1.08 Å) and IM7m (1.89 ± 1.10 Å) bound to mutated Gyr in the QRDR site, which implies the active interaction of these compounds with Gyr, indicating their underlying stability.

**Figure 3.**
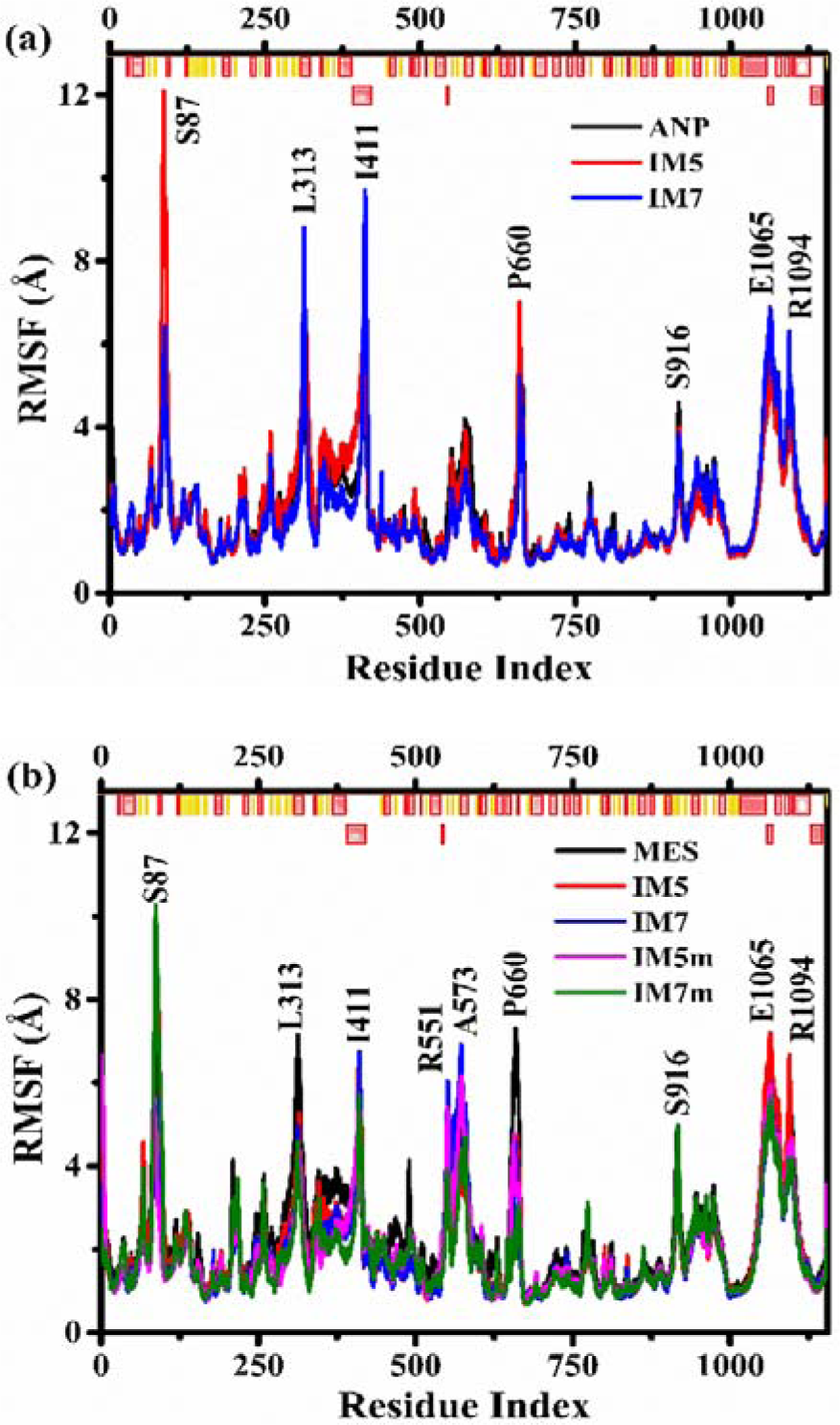
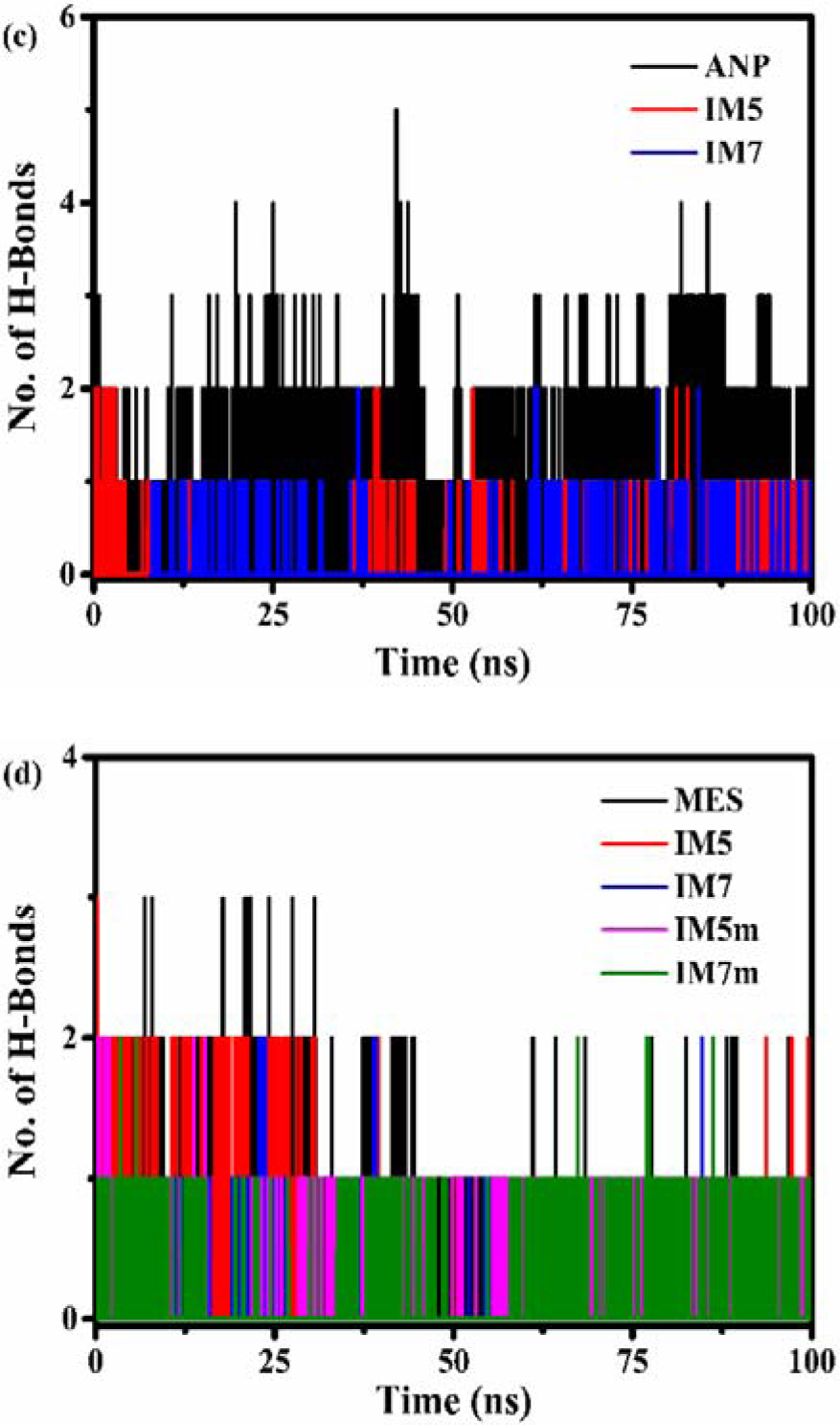
**(a)** Root-mean-square fluctuation of reference (ANP) and lead (IM5, IM7) compounds in the ATP site; **(b)** Root-mean-square fluctuation of reference (MES) and lead compounds in the QRDR (IM5, IM7) and mutated QRDR (IM7m, IM7m) site; **(c)** Number of hydrogen bonds plot for reference (ANP) and lead (IM5, IM7) compounds in the ATP site; **(d)** Number of hydrogen bonds plot for reference (MES) and lead compounds in the QRDR (IM5, IM7) and mutated QRDR (IM7m, IM7m) site. In **Figure (a)** and **(b)**, the top line colour codes represent helix, sheet, and loop regions in dark-red, yellow, and white, respectively.

The estimation of the stability of the interaction process was also done by analyzing the number of hydrogen bonds formed between receptor and ligand. The number of H-bonds formed between Gyr and the selected compounds from the MD simulations was calculated and provided in **Figure 3**. **Figures 3(c, d)** show a changing number of H-bonds between Gyr and the selected compounds, which implies that the ligands are changing conformation in the active site of the macromolecule during the simulation time. For the ATP binding site (**Figure 3(c)**), the Gyr-ANP complex had 1-5 H-bonds, while 1-2 H-bonds were there for Gyr-IM5 and Gyr-IM7 during the simulation. There are 1-3 H-bonds observed in the case of the QRDR site (**Figure 3(d)**) of Gyr for IM5 and the reference compound MES, while Gyr-IM7 had 1-2 H-bonds. The mutated (A743V and D747G) Gyr formed 1-2 H-bonds with IM5m and IM7m (**Figure 3(d)**) during any instant of the simulation. The residues of Gyr that are involved in the hydrogen bond interaction with IM5, IM7, and the reference compounds are shown in **Table S4**. The residues of Gyr that formed hydrogen bond interaction with the ligands in the docking study are also maintained during the simulation, which indicates to their underlying interaction stability. The presence of the H-bonds between Gyr and the lead compounds during the simulation implies the stability of these complexes

### 3.6. Principal component analysis

Usually, the functional activity of the macromolecules arises from the complex correlated motions among its atomic coordinates which are necessary for their specific functioning such as conformational changes to certain biological environments. Proteins perform many kinds of internal motions that are difficult to interpret. Principal component analysis is a useful tool that reduces a multidimensional data to a few principal components that efficiently describes the essential properties of the protein’s global motion. From the MD trajectories it helps determine the concerted motion of receptor and receptor-ligand complexes. Here the change in the protein trajectory is represented by the PCs (PC1 and PC2) that are the eigenvector values obtained from the covariance matrix during MD simulation [43]. First, the values of the eigenvectors were extracted from the simulation’s covariance matrix. Later the protein movements that are dominant were filtered from various trajectories. A 2D PCA was performed to analyze the Gyr dynamics in the absence and presence of ligands (**Figure 4**). **Figure 4(a)** shows the projection vectors of the principal components of reference (ANP) and lead (IM5 and IM7) compounds in the ATP site. The analysis of the individual PC plots of ANP (**Figure S6(a)**), IM5 (**Figure S6(b)**), and IM7 (**Figure S6(c)**) shows that the lead compound complexes maintained less flexible conformation as compared to their reference ligands for the ATP site. The superposed (**Figure 4(b)**) and individual PC plots of the ligands in the QRDR (**Figure S6(d-f)**) and mutated QRDR (**Figure S6(g, h)**) site also indicate for the reduced conformational flexibility of the lead molecules with respect to the reference ligand. The ligand IM7m in mutated Gyr (**Figure S6(j)**) showed the least flexibility compared to the compounds in the QRDR site, implying a stable complex formation of the complex.

**Figure 4.**
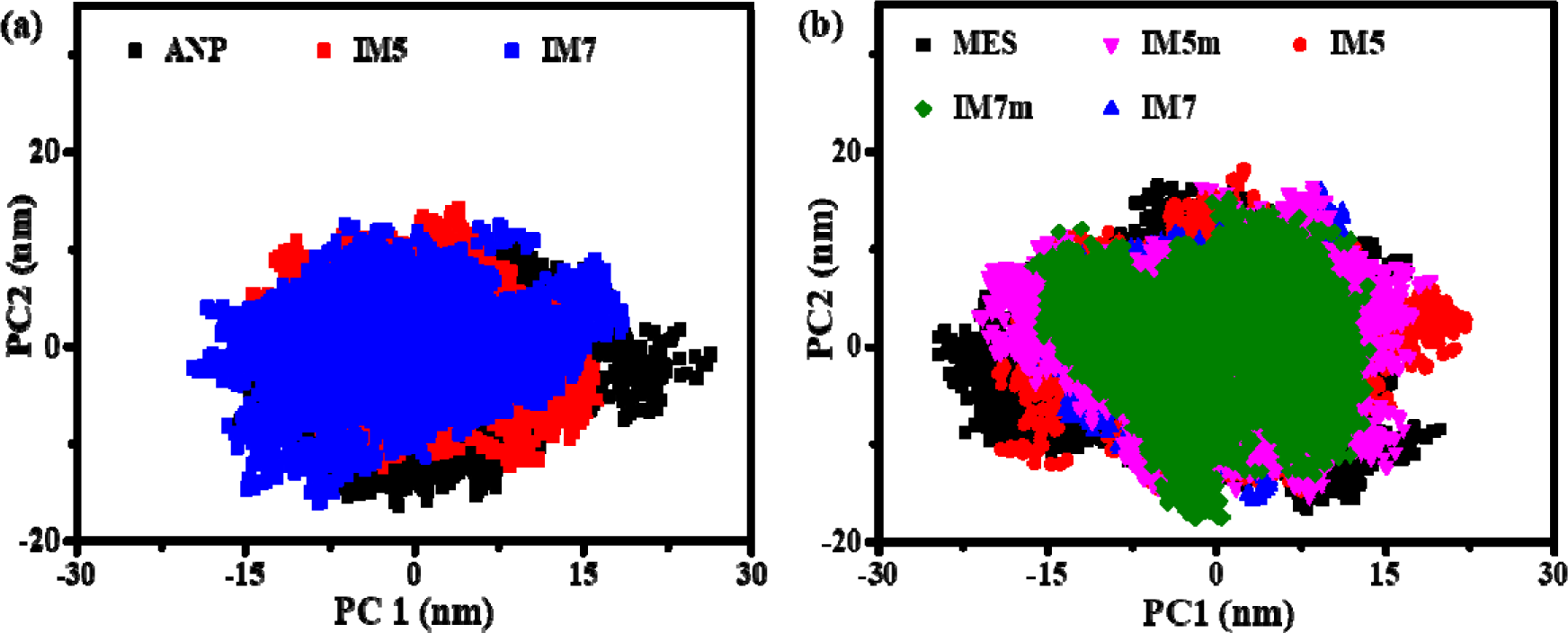
Principal components plot of compounds in the (a) ATP binding site, (b) QRDR and mutated QRDR site.

The Gibbs free energy landscape plots were made to analyze the energy minima of the Gyr-ligand complexes using the reaction coordinates obtained from the principal components (**Figure S7**). All the ligand-bound forms possess less free energy values than their corresponding reference ligands for both the ATP and QRDR sites, showing their stability and energetically favorable conformations. The low energy sampling regions (blue colour) are more for the IM5 and IM7 in all the cases than the reference compounds, suggesting these complexes’ thermodynamic favorability.

### 3.7. Binding free energy calculation

Calculating binding free energy is one of the preferred approaches to predict the binding affinity of the receptor-ligand complex. Estimation of binding free energy from the simulation trajectory can be used to scale the stability of the systems in terms of their nonbonded interactions. Here, we have used the MM/PBSA method to calculate the binding free energy of different systems considering 1000 snapshots from the trajectory equilibrium region. The binding free energies of the Gyr-ligand complexes are provided in **Table 1**. The negative affinity values from **Table 1** indicate that the interacting complexes are stable. The van der Waals interaction energy contributes majorly to the total affinity of the complexes. For all the complexes, van der Waals, electrostatics, and nonpolar interaction energies contributed actively except for the IM7 in the ATP binding site of Gyr, which is stabilized by polar, nonpolar, and van der Waals interaction energies.

**Table 1.** Contribution of different energy terms to MM/PBSA binding free energy of Gyr-ligand complexes as obtained from the MD trajectories.

| Binding Site | Molecule | $\Delta E_{\text{vdw}}$ | $\Delta E_{\text{ele}}$ | $\Delta G_{\text{pol}}$ | $\Delta G_{\text{nonpol}}$ | $\Delta G_{\text{Bind}}$ |
| --- | --- | --- | --- | --- | --- | --- |
| ATP | ANP | $-32.48 \pm 0.83$ | $-7.79 \pm 2.31$ | $18.57 \pm 0.71$ | $-4.08 \pm 0.14$ | $-25.78 \pm 4.14$ |
| | IM5 | $-19.31 \pm 1.37$ | $-2.28 \pm 0.36$ | $3.21 \pm 0.07$ | $-2.62 \pm 0.01$ | $-21.00 \pm 1.65$ |
| | IM7 | $-18.65 \pm 1.20$ | $3.80 \pm 0.49$ | $-1.30 \pm 0.62$ | $-2.68 \pm 0.01$ | $-18.83 \pm 1.62$ |
| QRDR | MES | $-21.53 \pm 0.45$ | $-3.55 \pm 0.12$ | $6.60 \pm 0.13$ | $-2.49 \pm 0.01$ | $-20.98 \pm 0.84$ |
| | IM5 | $-20.84 \pm 1.47$ | $-1.13 \pm 0.35$ | $3.34 \pm 0.09$ | $-2.69 \pm 0.00$ | $-21.32 \pm 1.71$ |
| | IM7 | $-29.72 \pm 1.23$ | $-6.86 \pm 0.64$ | $11.03 \pm 0.70$ | $-3.65 \pm 0.01$ | $-29.19 \pm 1.71$ |
| MQRDR | IM5 | $-30.21 \pm 1.40$ | $-3.98 \pm 0.50$ | $6.32 \pm 0.15$ | $-3.37 \pm 0.02$ | $-31.24 \pm 1.70$ |
| | IM7 | $-26.47 \pm 1.11$ | $-2.63 \pm 0.46$ | $5.37 \pm 0.45$ | $-3.25 \pm 0.01$ | $-26.97 \pm 1.50$ |

This research suggests that two molecules, IM5 and IM7, form stable complexes with the DNA Gyr enzyme of M. tuberculosis, making them promising candidates for treating tuberculosis infections. Docking studies showed that IM5 and IM7 were the most effective inhibitors among molecules screened from a large database (IMPPAT 2.0). They were effective against both the ATP and QRDR binding sites of DNA Gyr, even in mutated forms of the enzyme. Previous studies identified specific mutations in the Gyr enzyme of E. coli bacteria (S83 and D87) that cause resistance to quinolone antibiotics. These mutations correspond to residues A90 and D94 in M. tuberculosis (as shown in **Figure S2(a)**). These same residues are known to be crucial for the interaction between Gyr and quinolone drugs, and mutations in these areas can cause the bacteria to become roughly 10 times more resistant to the drugs [53]. MD simulations and MMPBSA analysis indicate that IM5 and IM7 form stable complexes with the QRDR site of both the normal and mutated forms of DNA Gyr. Docking analysis revealed that the hydrogen bonds formed between IM5/IM7 and specific residues of DNA gyrase are consistent with the findings from MD simulations (see **Table S4**). The fact that IM5 and IM7 form stable complexes in the DNA binding pocket (QRDR site) of Gyr, regardless of mutations in two key quinolone-interacting residues, suggests that these molecules could be effective inhibitors for both normal and mutated forms of the enzyme. This makes them attractive candidates for developing new drugs to combat tuberculosis.

## 4. Conclusion

This study employed in silico techniques, namely molecular docking, and MD simulation, to screen a natural molecule database (IMPPAT 2.0) for potential DNA gyrase inhibitors against *Mycobacterium tuberculosis* (MTB), including drug-resistant strains. Two molecules, IM5 and IM7, emerged as promising candidates based on their docking scores, drug-likeness, and predicted pharmacokinetic properties. The analysis of molecular electrostatic potential and docking interactions suggested favorable binding between these molecules and DNA gyrase. Furthermore, IM5 and IM7 exhibited ADMET-compliant properties, indicating their potential for further development. MD simulations supported the stability of the IM5-Gyr and IM7-Gyr complexes, with principal component analysis (PCA) and binding free energy calculations further suggesting their active role in complex formation. Based on these computational findings, IM5 and IM7 warrant further investigation as potential inhibitors of DNA gyrase for the treatment of MTB and its drug-resistant forms. However, *in vitro*, and *in vivo* studies are necessary to validate these results.

## Supporting information

https://drive.google.com/file/d/1_i0KpToJW2_RztFmKJ91SXXTpbVRuQkN/view?usp=drive_link

## Acknowledgements

The authors are thankful to the HPC facility, IIT Bombay for the computational resources.

## Authors’ Contributions

**Janmejaya Rout:** Conceptualization; Investigation; Formal analysis; Validation; Visualization; Writing - Original draft, Writing - Review & Editing; **Sayani Das:** Investigation; Validation; Writing - Review & Editing; **Sandip Kaledhonkar:** Conceptualization; Investigation; Validation; Funding acquisition; Project administration; Resources; Supervision; Writing-Review & Editing.

## Data availability statement

The data in support of the findings of this study are available within the article and its supplementary materials.

## Abbreviations

Gyr: Gyrase
MDR: Multidrug-resistant
XDR: Extensively-drug resistant
QRDR: Quinolone resistance-determining regions
IMPPAT 2.0: Indian Medicinal Plants, Phytochemistry and Therapeutics
PAINS: Pan assay interface
MEP: molecular electrostatic potential
ADMET: absorption, distribution, metabolism, excretion, and toxicity.

