## Supplementary material for "In Silico Identification of Potential Inhibitors of Mycobacterium tuberculosis DNA Gyrase from Phytoconstituents of Indian Medicinal Plants": https://drive.google.com/file/d/1_i0KpToJW2_RztFmKJ91SXXTpbVRuQkN/view?usp=drive_link

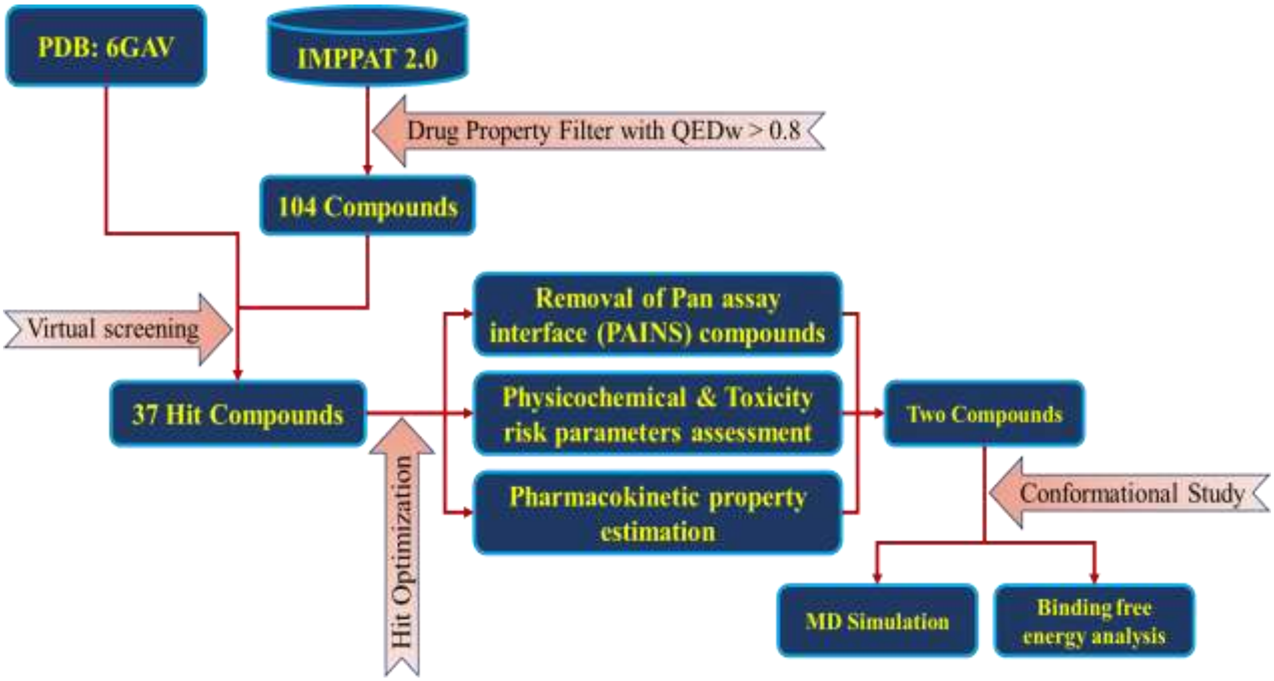


**Figure S1.** Flow chart of the study for the identification of novel DNA gyrase inhibitors. Potential compounds were screened from the IMPPAT 2.0 database resulting two lead compounds after molecular docking and pharmacokinetic filter whose conformational stability were monitored with the DNA gyrase by MD Simulation.

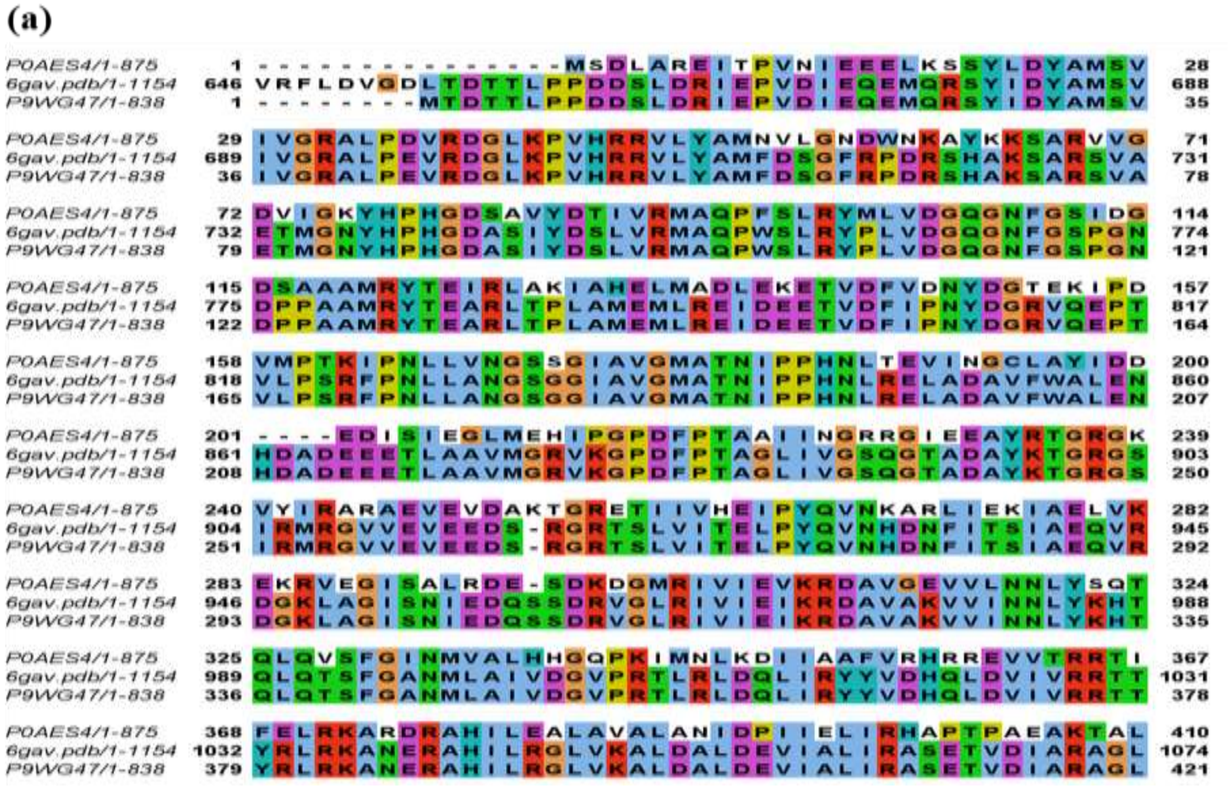

**(b)**

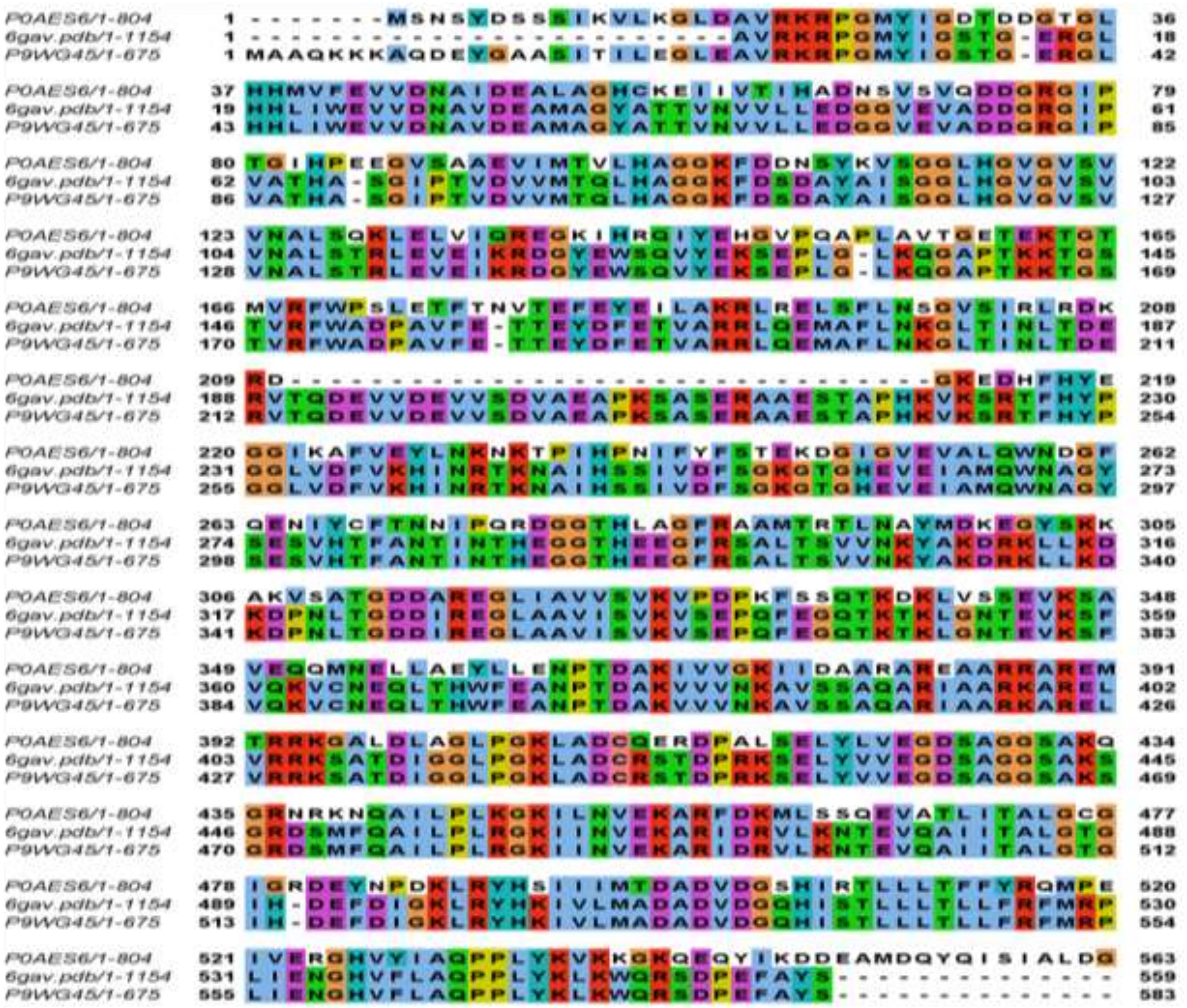

**Figure S2.** Sequence matching of *M. tuberculosis* (P9WG47) and *E. coli* (P0AES4) GyrA (a) and *M. tuberculosis* (P9WG45) and *E. coli* (P0AES6) GyrB (b) from UniProt with PDB 6GAV. The matched regions are coloured using clustal colour scheme in Jalview.

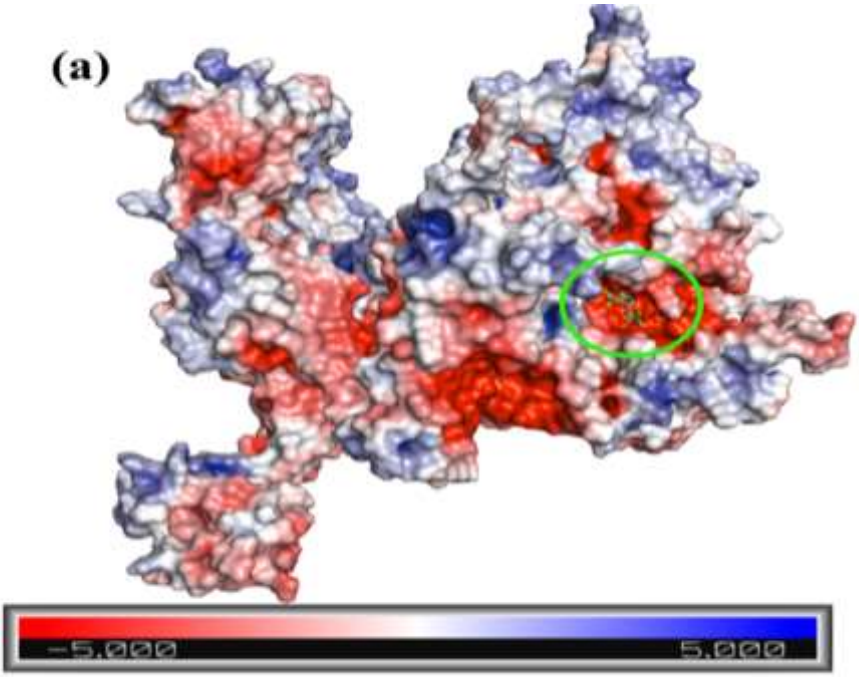
(a)
−5.000
5.000

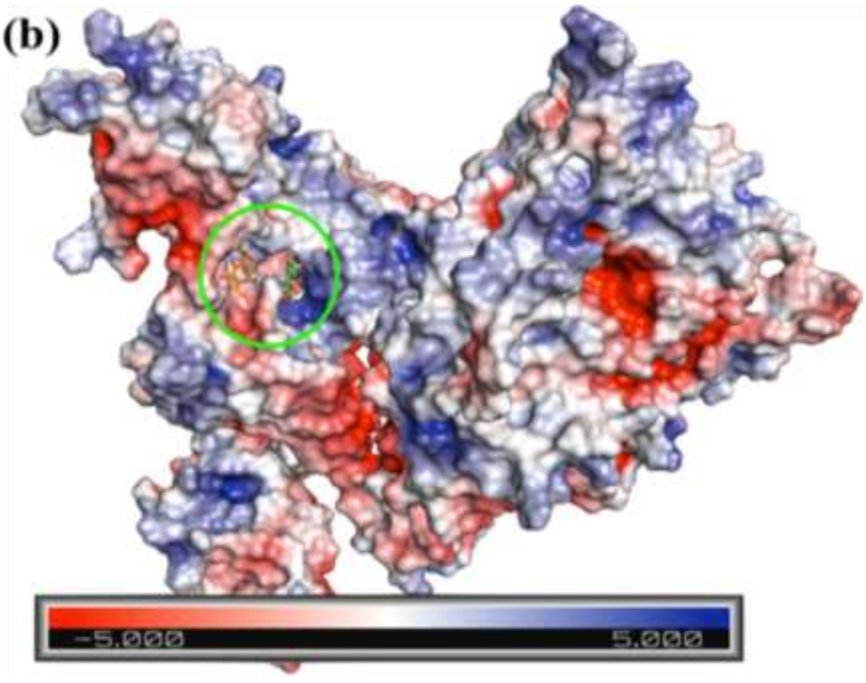
(b)
−5.000
5.000

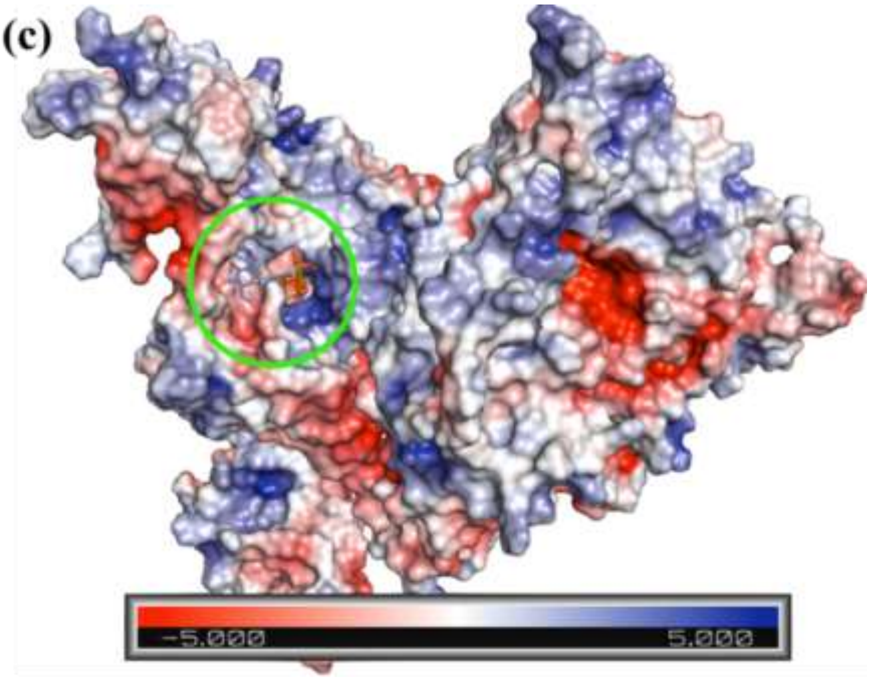


**Figure S3.** Molecular electrostatic potential energy surface plot of gyrase in presence of reference and lead compounds in the (a) ATP binding site, (b) QRDR site, and (c) mutated QRDR site. Here the electron deficient regions are in blue, neutral regions are in white, and electron rich regions are presented in red.

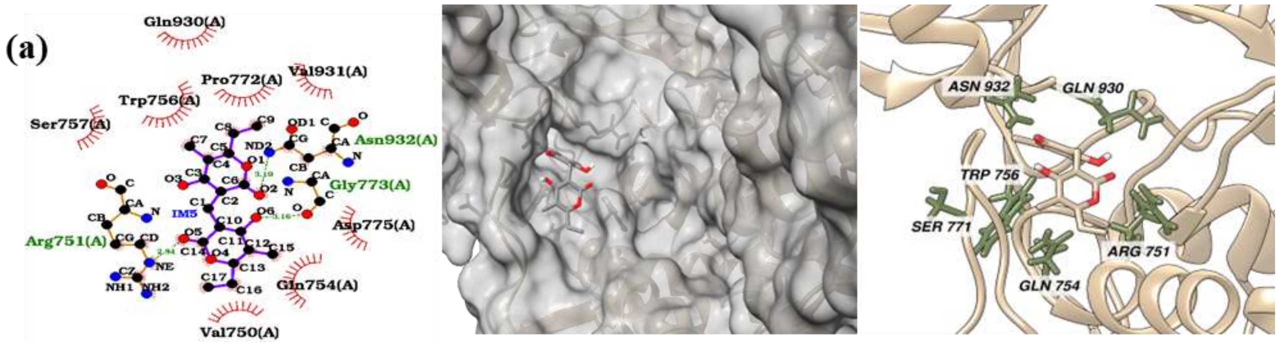

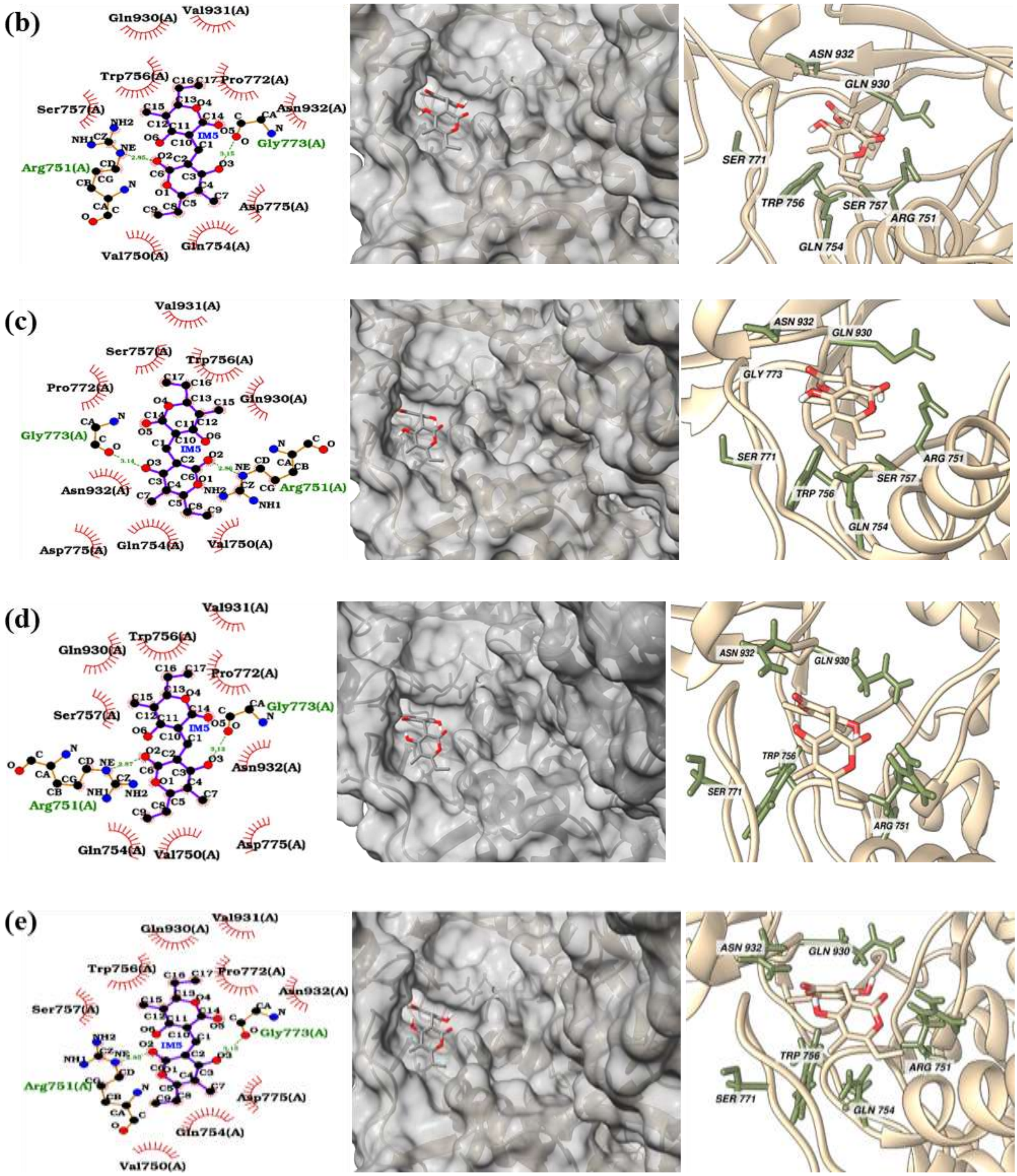
(b)
Gln930(A)
Val931(A)
Trp756(A)
Pro772(A)
Ser757(A)
Asn932(A)
Gly773(A)
Arg751(A)
Asp775(A)
Gln754(A)
Val750(A)
ASN 932
GLN 930
SER 771
TRP 756
SER 757
ARG 751
GLN 754
(c)
GLY 773
(d)
(e)

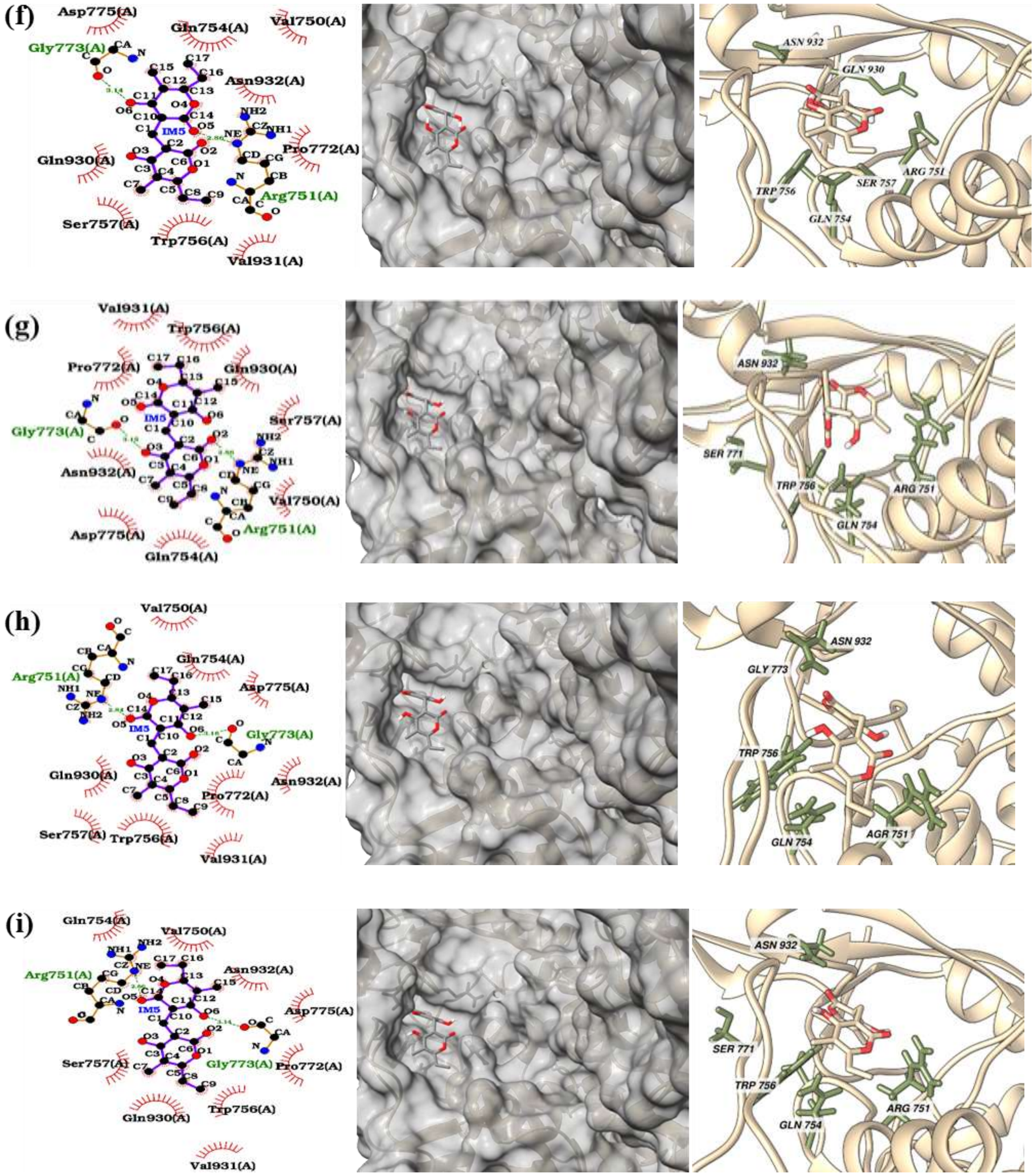
(f)
Asp775(A)
Gln754(A)
Val750(A)
Gly773(A)
Asn932(A)
IM5
Pro772(A)
Gln930(A)
Arg751(A)
Ser757(A)
Trp756(A)
Val931(A)
ASN 932
GLN 930
ARG 751
TRP 756
SER 757
GLN 754
(g)
Val931(A)
Trp756(A)
Pro772(A)
Gln930(A)
Ser757(A)
Gly773(A)
Asn932(A)
Val750(A)
Arg751(A)
Asp775(A)
Gln754(A)
ASN 932
SER 771
TRP 756
ARG 751
GLN 754
(h)
Val750(A)
Gln754(A)
Arg751(A)
Asp775(A)
Gly773(A)
Gln930(A)
Asn932(A)
Pro772(A)
Ser757(A)
Trp756(A)
Val931(A)
ASN 932
GLY 773
TRP 756
GLN 754
AGR 751
(i)
Gln754(A)
Val750(A)
Arg751(A)
Asn932(A)
Asp775(A)
Ser757(A)
Gly773(A)
Pro772(A)
Trp756(A)
Gln930(A)
Val931(A)
ASN 932
SER 771
TRP 756
GLN 754
ARG 751

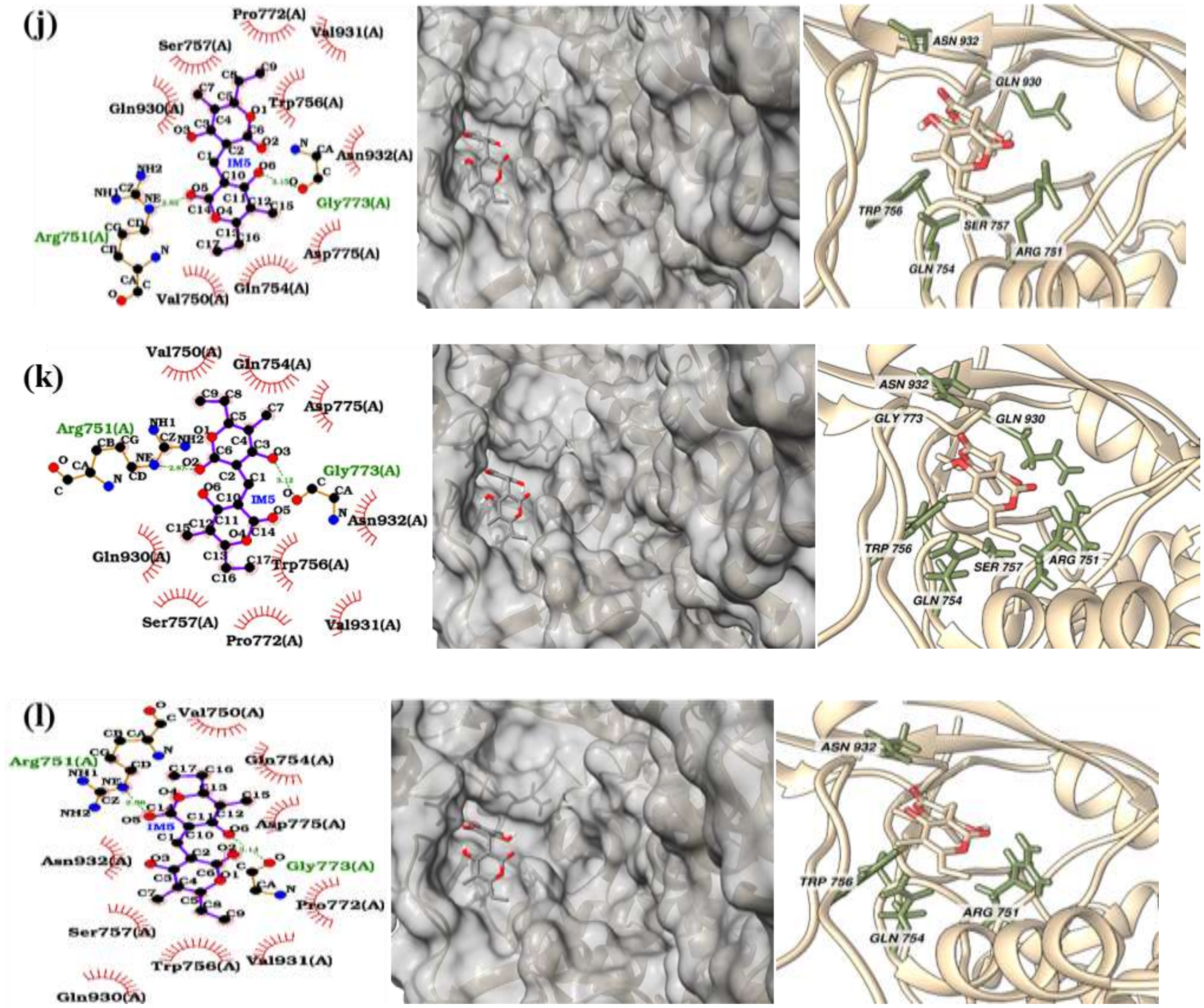


**Figure S4.** Respective 2D interaction diagram, surface, and 3D binding site view of the lowest energy docked pose of IM5 in the QRDR site of Gyr for the residual mutations **(a)** A743G and D747A, **(b)** A743G and D747G, **(c)** A473G and D747H, **(d)** A743G and D747N, **(e)** A743L and D9747A, **(f)** A743L and D747G, **(g)** A743L and D747H, **(h)** A743L and D747N, **(i)** A743V and D747A, **(j)** A743V and D747G, **(k)** A743V and D747H, **(l)** A743V and D747N.

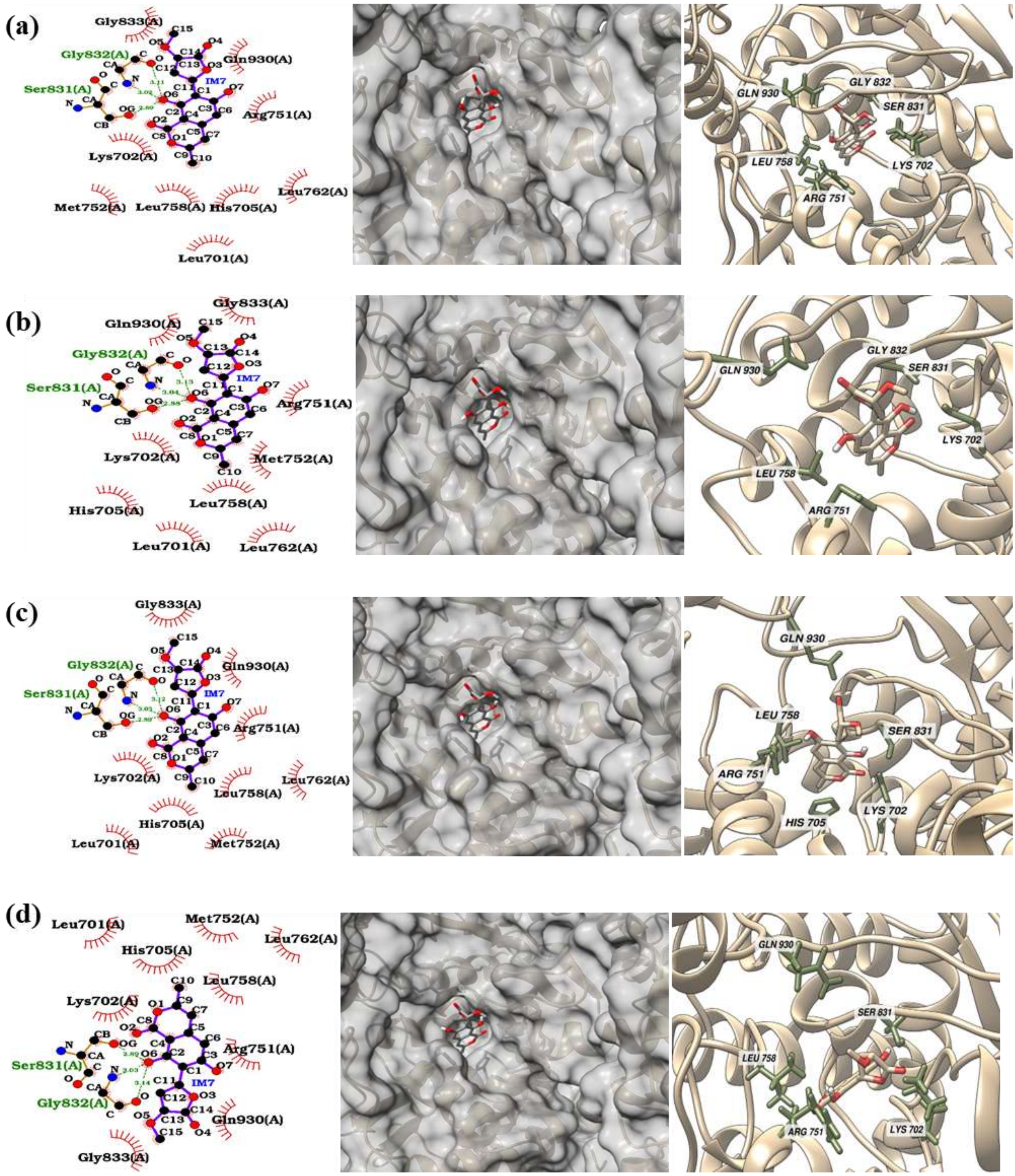
(a)
Gly833(A)
Gly832(A)
Ser831(A)
Gln930(A)
Arg751(A)
Lys702(A)
Met752(A)
Leu758(A)
His705(A)
Leu762(A)
Leu701(A)
IM7
GLN 930
GLY 832
SER 831
LEU 758
LYS 702
ARG 751
(b)
Gly833(A)
Gln930(A)
Gly832(A)
Ser831(A)
Arg751(A)
Lys702(A)
Met752(A)
His705(A)
Leu758(A)
Leu701(A)
Leu762(A)
IM7
GLN 930
GLY 832
SER 831
LYS 702
LEU 758
ARG 751
(c)
Gly833(A)
Gly832(A)
Gln930(A)
Ser831(A)
Arg751(A)
Lys702(A)
Leu762(A)
Leu758(A)
His705(A)
Leu701(A)
Met752(A)
IM7
GLN 930
LEU 758
SER 831
ARG 751
LYS 702
HIS 705
(d)
Leu701(A)
Met752(A)
Leu762(A)
His705(A)
Leu758(A)
Lys702(A)
Arg751(A)
Ser831(A)
Gly832(A)
Gln930(A)
Gly833(A)
IM7
GLN 930
SER 831
LEU 758
ARG 751
LYS 702

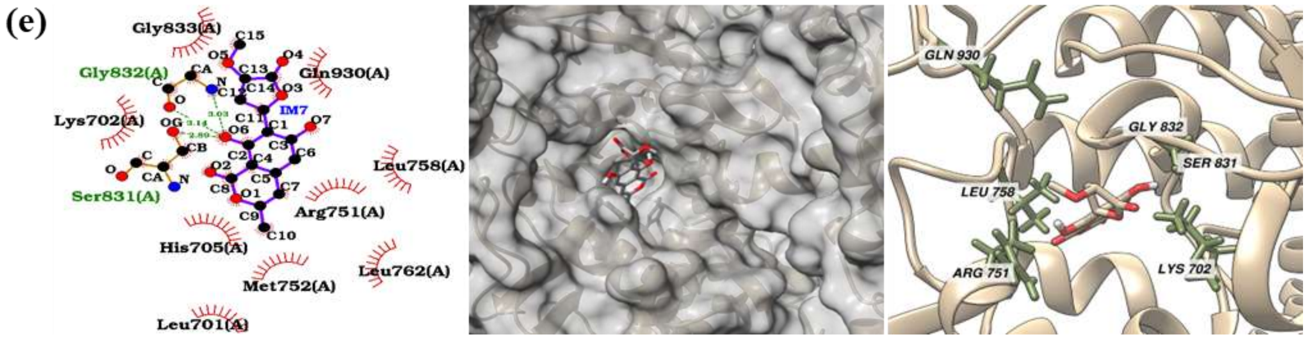
(e)
Gly833(A)
Gly832(A)
Gln930(A)
Lys702(A)
IM7
Leu758(A)
Ser831(A)
Arg751(A)
His705(A)
Leu762(A)
Met752(A)
Leu701(A)
GLN 930
GLY 832
SER 831
LEU 758
ARG 751
LYS 702

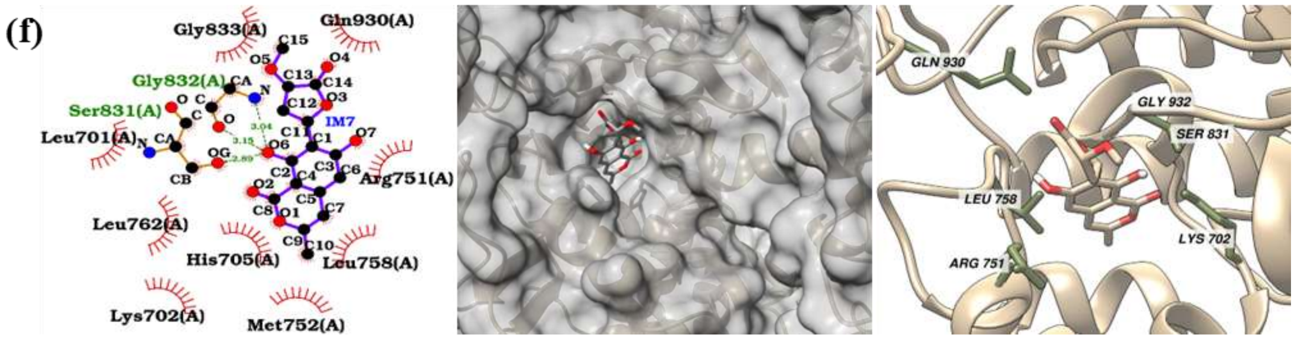
(f)
Gly833(A)
Gln930(A)
Gly832(A)
Ser831(A)
IM7
Leu701(A)
Arg751(A)
Leu762(A)
His705(A)
Leu758(A)
Lys702(A)
Met752(A)
GLN 930
GLY 932
SER 831
LEU 758
LYS 702
ARG 751

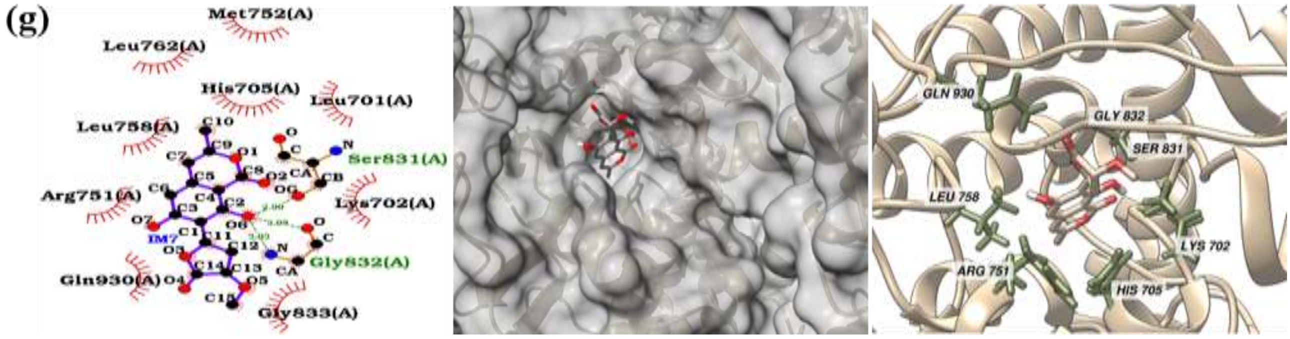
(g)
Met752(A)
Leu762(A)
His705(A)
Leu701(A)
Leu758(A)
Ser831(A)
Arg751(A)
Lys702(A)
IM7
Gly832(A)
Gln930(A)
Gly833(A)
GLN 930
GLY 832
SER 831
LEU 758
LYS 702
ARG 751
HIS 705

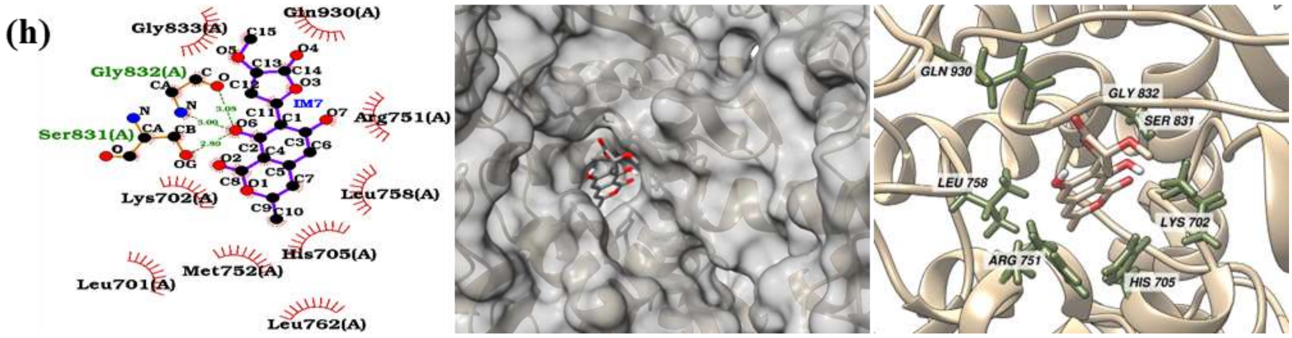
(h)
Gly833(A)
Gln930(A)
Gly832(A)
IM7
Arg751(A)
Ser831(A)
Lys702(A)
Leu758(A)
His705(A)
Met752(A)
Leu701(A)
Leu762(A)
GLN 930
GLY 832
SER 831
LEU 758
LYS 702
ARG 751
HIS 705

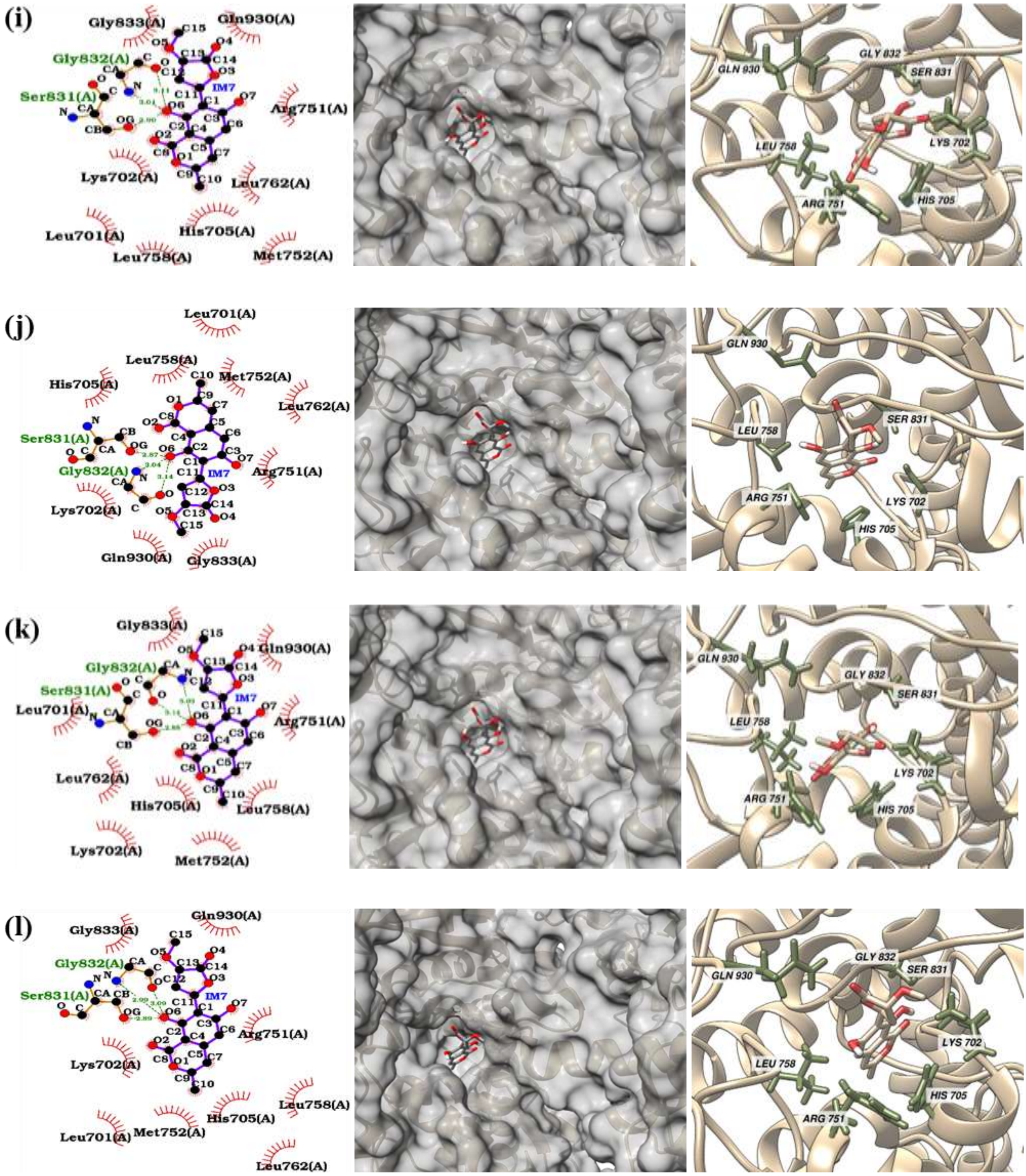


**Figure S5.** Respective 2D interaction diagram, surface, and 3D binding site view of the lowest energy docked pose of IM7 in the QRDR site of Gyr for the residual mutations **(a)** A743G and D747A, **(b)** A743G and D747G, **(c)** A473G and D747H, **(d)** A743G and D747N, **(e)** A743L and D9747A, **(f)** A743L and D747G, **(g)** A743L and D747H, **(h)** A743L and D747N, **(i)** A743V and D747A, **(j)** A743V and D747G, **(k)** A743V and D747H, **(l)** A743V and D747N.

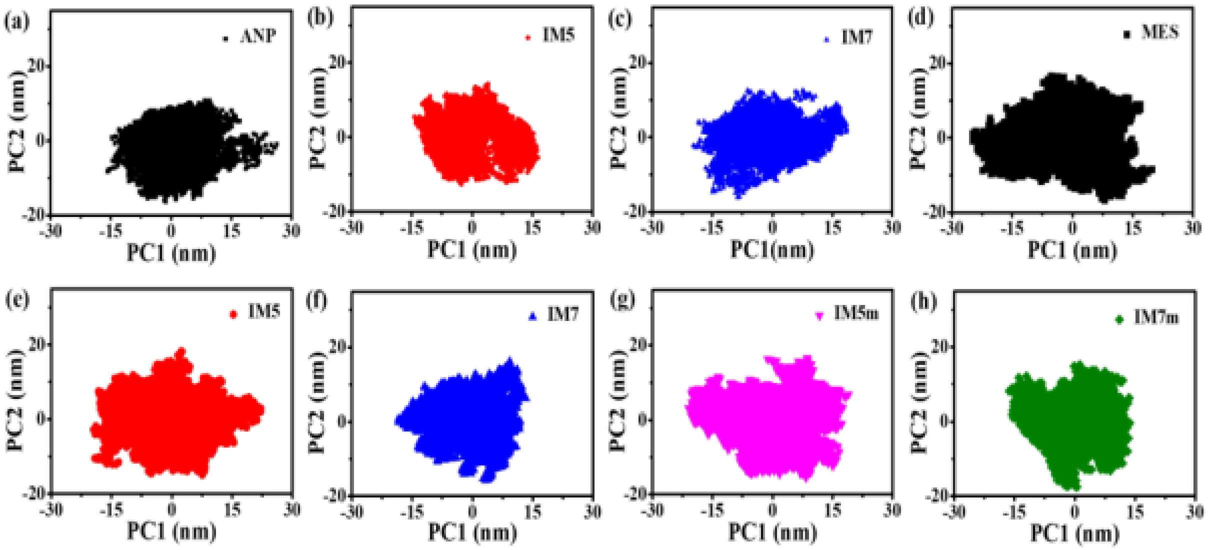


**Figure S6.** Principal components plot of Gyr in the presence of (a) ANP, (b) IM5, (c) IM7 in the ATP site. Principal components plot of Gyr in the presence of (d) MES, (e) IM5, (f) IM7 in the QRDR site and (g) IM5m; (h) IM7m in the mutated QRDR site.

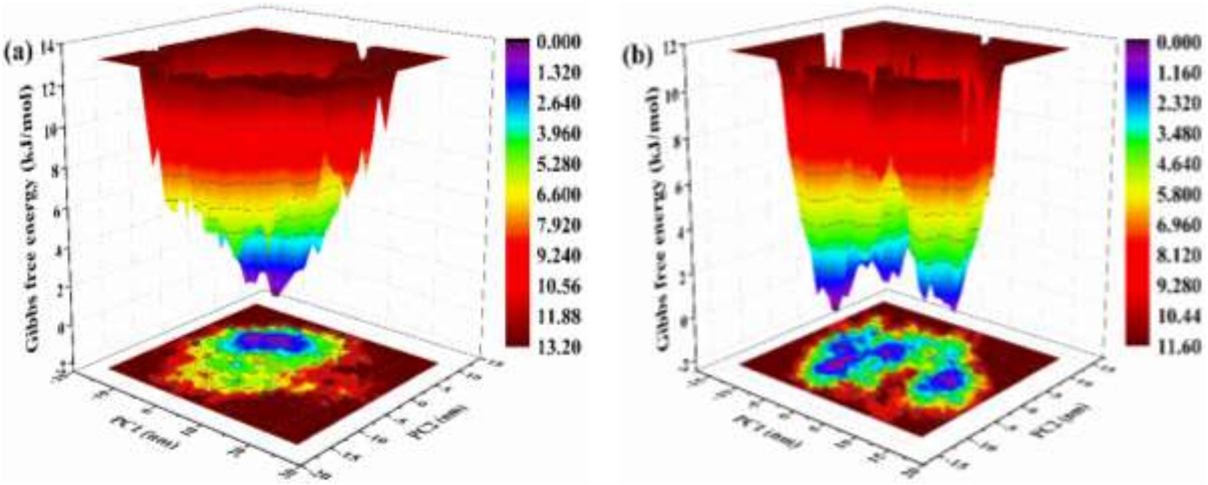

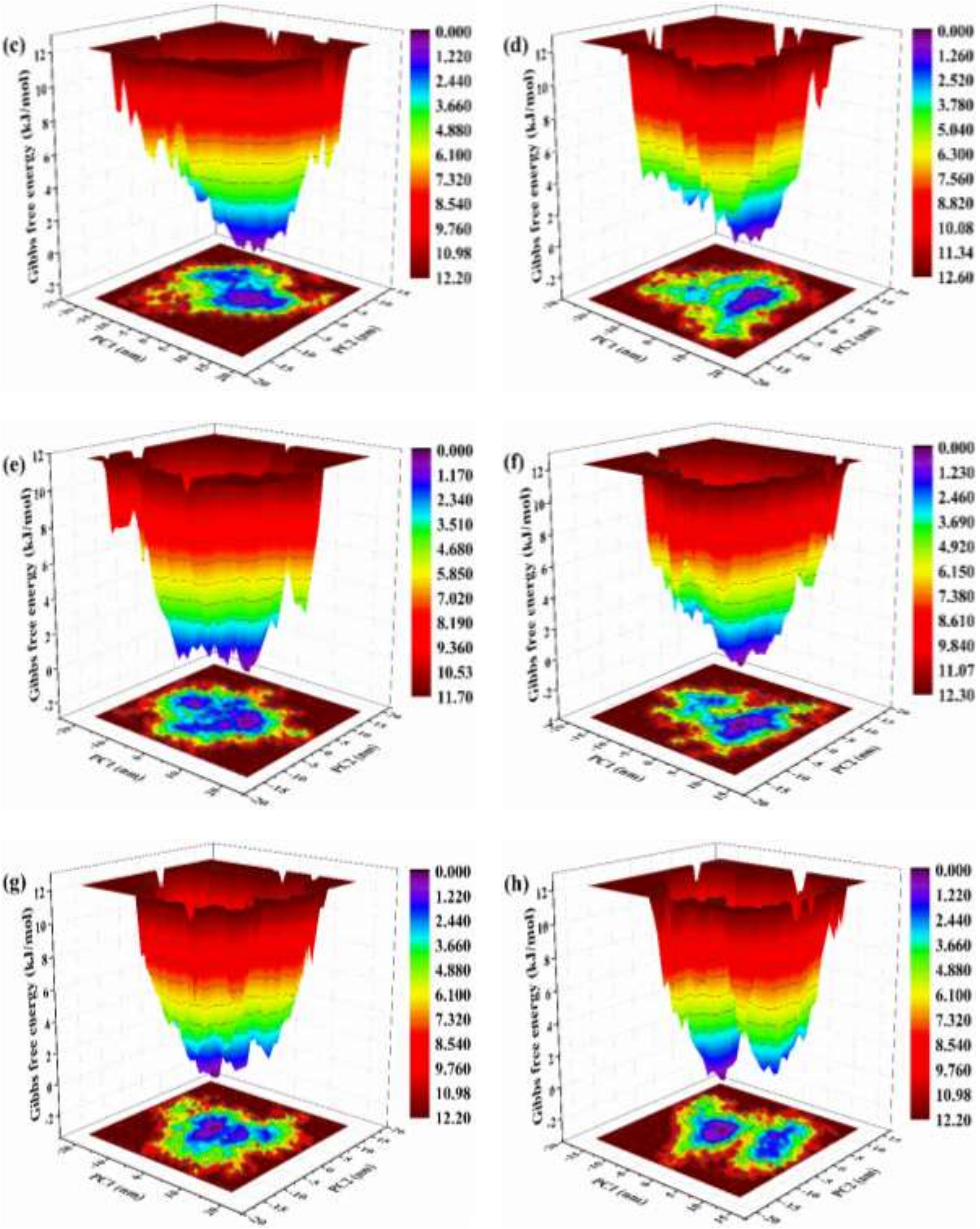


**Figure S7.** Free energy landscape plots of compounds as a function of projections of the MD trajectory onto the first (PC1) and second (PC2) eigenvectors, respectively in the ATP binding site (ANP **(a)**, IM5 **(b)**, and IM7 **(c)**), QRDR (MES **(d)**, IM5 **(e)**, and IM7 **(f)**), and mutated QRDR (IM5 **(g)** and IM7 **(h)**) site.

**Table S1.** Physicochemical and toxicity parameters of the reference and hit compounds (*: Unknown stereo configuration from OSIRIS property explorer during toxicity test). [MW: molecular weight; cLogP: Lipophilicity; HBA: Hydrogen bond acceptor; HBD: Hydrogen bond donor; TPSA: Topological polar surface area; RB: Rotatable bonds].

| Sl. No. | Molecule | Physicochemical Parameters | | | | | | | Toxicity Risk Parameters | |
|---|---|---|---|---|---|---|---|---|---|---|
| | | MW | cLogP | Solubility | HBA | HBD | TPSA | RB | Mutagenic | Tumorigenic |
| 1 | IMPHY006072 | 386.35 | 1.61 | -3.45 | 8 | 2 | 95.84 | 2 | No | No |
| 2 | IMPHY003597 | 370.35 | 2.24 | -3.69 | 7 | 1 | 75.61 | 2 | No | No |
| 3 | IMPHY003598 | 370.35 | 2.16 | -3.69 | 7 | 1 | 75.61 | 2 | No | No |
| 4 | IMPHY003954 | 386.35 | 1.67 | -2.96 | 8 | 2 | 95.84 | 2 | No | Yes |
| 5 | IMPHY004212 | 386.35 | 1.61 | -2.96 | 8 | 2 | 95.84 | 2 | No | Yes |
| 6 | IMPHY007572 | 328.32 | 2.7 | -4.25 | 6 | 2 | 85.22 | 2 | No | No |
| 7 | IMPHY011147 | 370.35 | 2.3 | -3.67 | 7 | 1 | 83.45 | 3 | No | No |
| 8 | IMPHY013734 | 386.35 | 1.39 | -2.77 | 8 | 2 | 95.84 | 2 | No | No |
| 9 | IMPHY012971 | 370.35 | 2.37 | -3.79 | 7 | 1 | 75.61 | 2 | No | No |
| 10 | IMPHY014962 | 370.35 | 2.37 | -3.79 | 7 | 1 | 75.61 | 2 | No | No |

| | | | | | | | | | | |
|---|---|---|---|---|---|---|---|---|---|---|
| 11 | IMPHY015035 | 370.35 | 2.07 | -3.35 | 7 | 1 | 75.61 | 2 | No | No |
| 12 | IMPHY010647 | 386.35 | 1.53 | -3.11 | 8 | 2 | 95.84 | 2 | No | No |
| 13 | IMPHY001390 | 358.39 | 0.77 | -2.26 | 6 | 2 | 88.1 | 3 | * | * |
| 14 | IMPHY003529 | 346.29 | 2.15 | -4.16 | 6 | 2 | 108.74 | 1 | No | No |
| 15 | IMPHY000536 | 348.43 | 2.72 | -3.53 | 5 | 2 | 79.9 | 3 | No | No |
| 16 | IMPHY000656 | 383.39 | 2.16 | -2.84 | 6 | 1 | 83.09 | 4 | No | No |
| 17 | IMPHY001304 | 370.4 | 2.53 | -4.14 | 6 | 2 | 77.38 | 1 | No | No |
| 18 | IMPHY001727 | 384.38 | 2.72 | -4.1 | 7 | 1 | 83.45 | 3 | No | No |
| 19 | IMPHY001907 | 383.39 | 1.98 | -3.66 | 7 | 1 | 77.46 | 2 | No | No |
| 20 | IMPHY004897 | 372.37 | 2.56 | -3.96 | 7 | 1 | 75.61 | 4 | No | No |
| 21 | IMPHY000144 | 370.35 | 2.19 | -3.42 | 7 | 1 | 83.45 | 4 | Yes | No |
| 22 | IMPHY001506 | 330.37 | 2.12 | -3.14 | 5 | 1 | 76.74 | 1 | No | No |
| 23 | IMPHY010864 | 330.29 | 2.05 | -3.6 | 7 | 2 | 86.61 | 1 | No | No |
| 24 | IMPHY011238 | 342.34 | 3.07 | -4.55 | 6 | 2 | 85.22 | 2 | No | No |
| 25 | IMPHY012973 | 301.34 | 0.94 | -2.14 | 5 | 2 | 76.38 | 2 | No | No |
| 26 | IMPHY014727 | 385.41 | 1.64 | -3.15 | 7 | 2 | 80.62 | 2 | No | No |

| | | | | | | | | | | |
|---|---|---|---|---|---|---|---|---|---|---|
| 27 | IMPHY000769 | 330.29 | 2.11 | -3.83 | 7 | 2 | 94.45 | 2 | No | No |
| 28 | IMPHY003137 | 370.35 | 2.1 | -3.35 | 7 | 1 | 75.61 | 2 | No | No |
| 29 | IMPHY005869 (IM5) | 320.34 | 2.53 | -3.05 | 6 | 2 | 100.88 | 4 | No | No |
| 30 | IMPHY010079 | 385.41 | 1.64 | -3.15 | 7 | 2 | 80.62 | 2 | No | No |
| 31 | IMPHY004724 | 344.32 | 2.17 | -3.8 | 7 | 1 | 75.61 | 2 | No | No |
| 32 | IMPHY006056 | 356.37 | 2.59 | -3.95 | 6 | 2 | 85.22 | 3 | No | No |
| 33 | IMPHY007403 (IM7) | 306.27 | 1.53 | -3.14 | 7 | 2 | 106.2 | 2 | No | No |
| 34 | IMPHY012646 | 371.38 | 2 | -3.43 | 7 | 2 | 78.41 | 2 | No | No |
| 35 | IMPHY013400 | 344.32 | 2.4 | -3.8 | 7 | 1 | 75.61 | 2 | No | No |
| 36 | IMPHY014004 | 374.43 | 1.22 | -2.71 | 6 | 2 | 80.26 | 3 | * | * |
| 37 | ANP | 506.2 | -4.48 | 1.11 | 16 | 8 | 311.36 | 8 | No | No |
| 38 | MES | 195.24 | -0.89 | 1.35 | 5 | 1 | 75.22 | 3 | No | No |

**Table S2.** Molecular docking and pharmacokinetic properties of the reference and hit compounds.

| Sl. No. | Molecule | Binding Affinity (kcal/mol) | | | Pharmacokinetic Parameters | | | | | | |
|---|---|---|---|---|---|---|---|---|---|---|---|
| | | ATP | QRDR | MQRDR | GI absorption | BBB Permeant | Pgp Substrate | CYP/I | Log Kp (cm/s) | Bioavailability score | Synthetic accessibility |
| 1 | IMPHY006072 | -8.6 | -9.4 | -9.2 | High | No | No | Yes | -7.49 | 0.55 | 4.5 |
| 2 | IMPHY003597 | -8.5 | -9.4 | -9.4 | High | No | No | Yes | -7.03 | 0.55 | 4.36 |
| 3 | IMPHY003598 | -8.5 | -9.4 | -9.2 | High | No | No | Yes | -7.03 | 0.55 | 4.36 |
| 4 | IMPHY007572 | -8.4 | -8.4 | -8.4 | High | No | No | Yes | -5.9 | 0.55 | 3.5 |
| 5 | IMPHY011147 | -8.2 | -8.2 | -8.3 | High | No | No | Yes | -6.98 | 0.55 | 4.17 |
| 6 | IMPHY013734 | -8.2 | -9.2 | -9.1 | High | No | No | Yes | -8.26 | 0.55 | 4.27 |
| 7 | IMPHY012971 | -8.1 | -9.0 | -8.7 | High | Yes | No | Yes | -6.91 | 0.55 | 4.31 |
| 8 | IMPHY014962 | -8.1 | -9.0 | -8.7 | High | Yes | No | Yes | -6.91 | 0.55 | 4.31 |
| 9 | IMPHY015035 | -8.1 | -8.8 | -8.8 | High | No | No | Yes | -7.41 | 0.55 | 4.22 |
| 10 | IMPHY010647 | -8.0 | -9.2 | -8.9 | High | No | No | Yes | -7.88 | 0.55 | 4.46 |
| 11 | IMPHY001390 | -7.8 | -7.8 | -7.8 | High | No | Yes | Yes | -8.23 | 0.55 | 5.48 |

| | | | | | | | | | | | |
|---|---|---|---|---|---|---|---|---|---|---|---|
| 12 | IMPHY003529 | -7.8 | -9.6 | -9.6 | High | No | No | Yes | -6.28 | 0.55 | 3.14 |
| 13 | IMPHY000536 | -7.7 | -7.7 | -7.7 | High | No | Yes | Yes | -6.65 | 0.55 | 4.95 |
| 14 | IMPHY000656 | -7.7 | -8.3 | -8.3 | High | No | Yes | Yes | -8.03 | 0.55 | 3.84 |
| 15 | IMPHY001304 | -7.7 | -8.7 | -8.6 | High | No | Yes | Yes | -6.59 | 0.55 | 4.36 |
| 16 | IMPHY001727 | -7.7 | -7.9 | -7.9 | High | No | No | Yes | -6.66 | 0.55 | 4.2 |
| 17 | IMPHY001907 | -7.7 | -8.3 | -8.2 | High | No | Yes | Yes | -7.22 | 0.55 | 4.54 |
| 18 | IMPHY004897 | -7.7 | -8.2 | -8.3 | High | Yes | No | Yes | -6.6 | 0.55 | 4.5 |
| 19 | IMPHY001506 | -7.6 | -7.7 | -7.5 | High | Yes | Yes | No | -7.01 | 0.55 | 4.91 |
| 20 | IMPHY010864 | -7.6 | -8.0 | -8.0 | High | No | Yes | Yes | -6.73 | 0.55 | 3.94 |
| 21 | IMPHY011238 | -7.6 | -8.3 | -8.3 | High | No | No | Yes | -5.73 | 0.55 | 3.61 |
| 22 | IMPHY012973 | -7.5 | -8.3 | -7.9 | High | No | Yes | No | -7.88 | 0.55 | 3.82 |
| 23 | IMPHY014727 | -7.5 | -7.9 | -7.7 | High | No | Yes | Yes | -7.82 | 0.55 | 4.76 |
| 24 | IMPHY000769 | -7.4 | -8.2 | -8.2 | High | No | Yes | Yes | -6.4 | 0.55 | 3.64 |
| 25 | IMPHY003137 | -7.4 | -8.8 | -8.6 | High | No | No | Yes | -7.41 | 0.55 | 4.22 |
| 26 | IMPHY005869 (IM5) | -7.4 | -7.4 | -7.3 | High | No | No | No | -7.01 | 0.55 | 3.81 |

| | | | | | | | | | | | |
|---|---|---|---|---|---|---|---|---|---|---|---|
| 27 | IMPHY010079 | -7.4 | -8.0 | -7.9 | High | No | Yes | Yes | -7.82 | 0.55 | 4.76 |
| 28 | IMPHY004724 | -7.3 | -7.8 | -7.9 | High | No | No | Yes | -6.59 | 0.55 | 4.06 |
| 29 | IMPHY006056 | -7.3 | -8.0 | -7.9 | High | No | Yes | Yes | -6.49 | 0.55 | 3.85 |
| 30 | IMPHY007403 (IM7) | -7.3 | -7.5 | -7.6 | High | No | No | No | -6.82 | 0.55 | 3.78 |
| 31 | IMPHY012646 | -7.3 | -8.2 | -8.2 | High | No | Yes | Yes | -7.34 | 0.55 | 4.41 |
| 32 | IMPHY013400 | -7.3 | -7.7 | -7.8 | High | Yes | No | Yes | -6.59 | 0.55 | 4.03 |
| 33 | IMPHY014004 | -7.3 | -8.3 | -8.3 | High | No | Yes | Yes | -7.93 | 0.55 | 5.81 |
| 34 | ANP | -7.2 | | | Low | No | Yes | No | -13.64 | 0.11 | 5.23 |
| 35 | MES | | -5.0 | -5.1 | High | No | No | No | -9.98 | 0.55 | 2.46 |

**Table S3.** Binding affinity and hydrogen bond analysis of IM5 and IM7 for the residual mutations made in the QRDR region as seen from the molecular docking study.

| Sl. No. | Mutation | Binding Affinity (kcal/mol) | | H-Bonding Residues | |
|---|---|---|---|---|---|
| | | IM5 | IM7 | IM5 | IM7 |
| 1 | A743G and D747A | -7.3 | -7.6 | R751, G773, N932 | S831, G832 |
| 2 | A743G and D747G | -7.3 | -7.6 | R751, G773 | S831, G832 |
| 3 | A473G and D747H | -7.3 | -7.6 | R751, G773 | S831, G832 |
| 4 | A743G and D747N | -7.3 | -7.6 | R751, G773 | S831, G832 |
| 5 | A743L and D9747A | -7.3 | -7.5 | R751, G773 | S831, G832 |
| 6 | A743L and D747G | -7.3 | -7.6 | R751, G773 | S831, G832 |
| 7 | A743L and D747H | -7.4 | -7.6 | R751, G773 | S831, G832 |
| 8 | A743L and D747N | -7.3 | -7.5 | R751, G773 | S831, G832 |
| 9 | A743V and D747A | -7.3 | -7.6 | R751, G773 | S831, G832 |
| 10 | A743V and D747G | -7.3 | -7.6 | R751, G773 | S831, G832 |
| 11 | A743V and D747H | -7.4 | -7.6 | R751, G773 | S831, G832 |
| 12 | A743V and D747N | -7.3 | -7.5 | R751, G773 | S831, G832 |

**Table S4.** The residues forming hydrogen bond interactions with IM5, IM7, and the reference compounds (ANP and MES) in the ATP binding site, QRDR and mutated QRDR sites of DNA gyrase as seen from MD simulation.

| Binding Site | IM5 | IM7 | MES | ANP |
|---|---|---|---|---|
| ATP Site | A1, R5, R16, H20, E24, D27, S210, R212, S216, T217, | M8, Y9, H20, H80, G94, H97, G98, K348, S423, T424, | | R3, K4, G7, M8, Y9, I10, G11, S12, R16, H20, E24, |

| | | | | |
|---|---|---|---|---|
| | K348, D615, R647, L649, E801, T1031, R1035, Q1128 | K428, D425, D448, H536, D1141, R1142 | | N285, H287, Q346, T347, K348, T349, T613, D615, D666, R667, E669 |
| QRDR site | R751, Q754, W756, S757, S771, G773, N774, D775, N932, N935, T938, R945, I952, S953, N954, I955, D957 | K702, H705, S748, R751, S757, N829, G830, G832, Y929, Q930 | K702, H705, R751, S757, N825, N829, S831, G832, Y929, Q930 | |
| Mutated QRDR site | R751, Q754, W756, S771, M780, N932 | R692, K702, H705, S748, R751, S757, N829, G832, I834, N841, Y929, Q930, N932 | | |
